# Oxygen levels at the time of activation determine T cell persistence and immunotherapeutic efficacy

**DOI:** 10.1101/2022.11.25.517976

**Authors:** Pedro P. Cunha, Eleanor Minogue, Lena C. M. Krause, Rita M. Hess, David Bargiela, Brennan J. Wadsworth, Laura Barbieri, Carolin Brombach, Iosifina P. Foskolou, Ivan Bogeski, Pedro Veliça, Randall S. Johnson

**Affiliations:** Department of Physiology, Development and Neuroscience, University of Cambridge, UK; Molecular Physiology, Institute of Cardiovascular Physiology, University Medical Center, Georg-August-University, Göttingen, Germany; Cancer Research UK, Cambridge Institute, University of Cambridge, UK; Department of Cell and Molecular Biology, Karolinska Institute, Sweden

**Keywords:** Immunology, CD8^+^ T cell, hypoxia, HIF, immunotherapy, CAR-T cells

## Abstract

**Summary:** Oxygenation levels are a determinative factor in T cell function. Here we describe that the oxygen tensions sensed by T cells at the moment of activation act to persistently modulate both differentiation and function. We found that in a protocol of CAR-T cell generation, 24 hours of low oxygen levels during initial CD8^+^ T cell priming is sufficient to enhance antitumour cytotoxicity in a preclinical model. This is the case even when CAR-T cells are subsequently cultured under high oxygen tensions prior to adoptive transfer. Increased hypoxia inducible transcription factor (HIF) expression was able to alter T cell fate in a similar manner to exposure to low oxygen tensions; however, only a controlled or temporary increase in HIF signalling was able to consistently improve cytotoxic function of T cells. These data show that oxygenation levels during and immediately after T cell activation play an essential role in regulating T cell function.

## Introduction

Cytotoxic T cells are often faced with oxygen-poor microenvironments (Labani-Motlagh et al., 2020), and in this context, tissue oxygen scarcity (hypoxia) is known to contribute to immune tolerance in tumours (Noman et al., 2015). For example, tumour hypoxia recruits immunosuppressive regulatory T cells, which can inhibit CD8^+^ T cell function (Facciabene et al., 2011). Hypoxia is also known to decrease expression of MHC class I molecules in tumour cells themselves, which in turn hampers T cell priming and cytotoxic function (Sethumadhavan et al., 2017). Tumour hypoxia can also have direct inhibitory effects on CD8^+^ T cells by increasing the expression of checkpoint inhibitor receptors (Bannoud et al., 2021; Noman et al., 2015) and by causing mitochondrial dysfunction (Liu et al., 2020; Scharping et al., 2021). However, T cells infiltrate a wide variety of tissues throughout their progression to maturation and final differentiation; they therefore need to function within a wide range of oxygen partial pressures, including oxygenation levels bordering on anoxia in some lymphoid organs (Caldwell et al., 2001). Thus, oxygen sensing mechanisms in T cells must be finely tuned for proper functioning in various tissue microenvironments.

In both preclinical studies and in clinical manufacturing, CD8^+^ T cells are typically cultured at atmospheric sea level O_2_ conditions (approximately 20.5%); however, reports have shown that physiologically relevant oxygen tensions (corresponding to 1%-5% O_2_) can profoundly shift T cell differentiation and function (Atkuri et al., 2007; Caldwell et al., 2001; Gropper et al., 2017; Larbi et al., 2010; Loeffler et al., 1990; Naldini et al., 1997; Palazon et al., 2017; Ross et al., 2021; Veliça et al., 2021). Lower oxygen levels alter CD8^+^ T cell effector differentiation in a process that requires hypoxia-inducible factors (HIFs), the key transcriptional regulators of cellular responses to low oxygen (Finlay et al., 2012; Palazon et al., 2017).

Culturing CD8^+^ T cells *ex vivo* at 1 %O_2_ has been found to dramatically reduce T cell proliferation, but increase anti-tumour activity, upon transfer into tumour-bearing mice (Gropper et al., 2017). Reduced oxygenation and increased HIF-α levels can thus both impair and enhance CD8^+^ T cell cytotoxicity. We hypothesised that this might be related to the timing of hypoxic stress during T cell activation and differentiation.

In this study, by combining the use of low oxygen conditioning with pharmacological inhibition of negative HIF regulators, we show that hypoxia has an unexpected role at the time point of activation, and for a short period thereafter. The effects of a short hypoxic conditioning are highly persistent, eliciting both *ex vivo* and *in vivo* increases in effector T cell differentiation and antitumour functioning, and demonstrate the critical role oxygen plays in the earliest stages of T cell activation.

## Results

### Hypoxia and increased HIF signalling drive CD8^+^ T cell effector differentiation

To assess the effect of different oxygen tensions in T cells, we first isolated polyclonal CD8^+^ T cells from C57BL/6J mice that were activated with anti-CD3 and anti-CD28 dynabeads for 3 days in ambient oxygen (21%), and 5% or 1% O_2_. Low oxygen levels increased effector differentiation of T cells, upregulating molecules such as the IL-2 receptor alpha subunit CD25, the transcription factors EOMES, T-bet and TOX, and the effector proteins granzyme B (GZMB) and interferon-γ (IFN-γ) (**Figure 1A**). T cells cultured at 1% O_2_ have increased expression of inhibitory receptors and impaired clonal expansion (**Figure 1B and S1A**). The HIF transcription factors are one of the primary mechanisms by which T cells respond to hypoxia and are critical to T cell function (Palazon et al., 2017). Contrary to many cell types, an increase in HIF-1α mRNA and protein levels can occur in T cells even at high oxygen levels, following stimulation of the T cell receptor (TCR) (Finlay et al., 2012; Lukashev et al., 2006; Makino et al., 2003; Nakamura et al., 2005; Wang et al., 2011). We first wished to determine whether the kinetics of HIF-1α stabilisation in *ex vivo* activated CD8^+^ T cells differs, depending on surrounding oxygen tensions (**Figure 1C**). In all oxygen tensions tested, HIF-1α protein expression increased after T cell activation and peaked within 24 hours. Exposure to 1% O_2_ resulted in the greatest accumulation of HIF-1α during the 3 days of culture (**Figure 1C**). HIF-2α protein was also increased in T cells following activation, but was mostly found in the cytoplasmic protein fraction (**Figure S1B**), as previously seen (Park et al., 2003). This is consistent with data showing that deletion of HIF-1α, but not HIF-2α, abrogates hypoxia-driven increases in GZMB (**Figure S1C**) and other activation markers, and impairs antitumour T cell function (Palazon et al., 2017).

**Figure 1.**
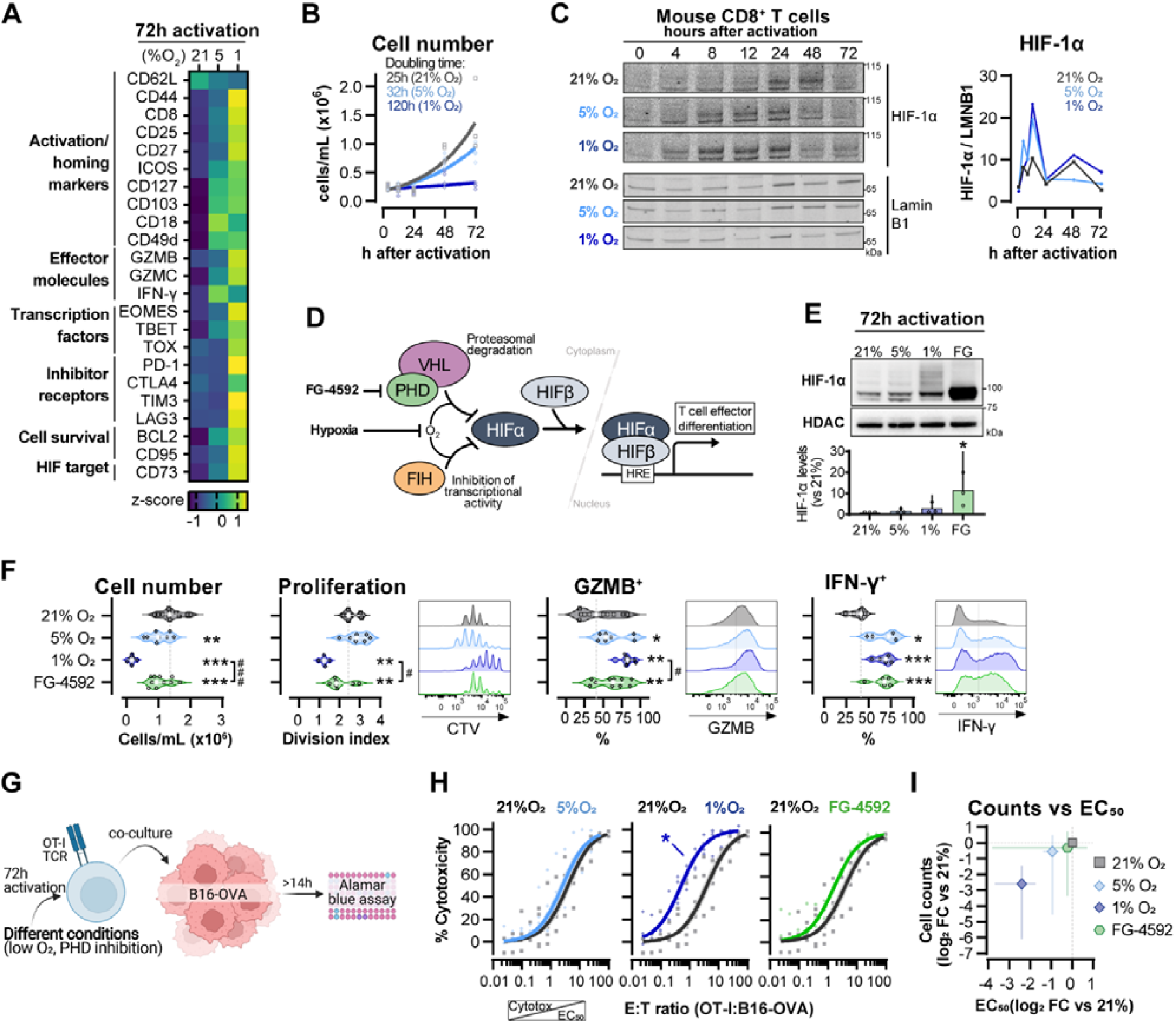
Hypoxia and inhibition of negative HIF signalling in CD8^+^ T cell effector differentiation. **(A)** Heat map illustrating expression of markers of differentiation as median fluorescence intensity (MFI) determined by flow cytometry 72h after activation of naive CD8^+^ T cells in 21%, 5% or 1% O_2_. Viridis was used as colour scale and rows represent averaged z-scores; n=4-29. **(B)** Number of CD8^+^ T cells following activation in 21%, 5% or 1% O_2_ shown in cells/mL and determined with automated cell counter. Exponential growth curve fit (represented by thick lines) was used to calculate doubling time in hours of T cells in each oxygen tension; n=4. **(C)** HIF-1α protein expression in nuclear extracts from CD8^+^ T cells activated in 21% (grey), 5% (light blue) or 1% (dark blue) O_2_. Representative immunoblots (left) and HIF-1α protein signal normalized to lamin B1 (right); n=1. **(D)** Representation of inhibition of negative regulators of HIFα with hypoxia and chemical inhibition of PHD with FG-4592. **(E)** HIF-1α protein expression in nuclear extracts from OT-I CD8^+^ T cells activated for 72h. Conditions analysed: wild-type (WT) cells activated in 21%, 5% or 1% O_2_; WT cells activated in 21% O_2_ and treated with 50 μM FG-4592. Representative immunoblot (top) and HIF-1α levels relative to 21% after normalization to HDAC or Histone 3 loading controls (bottom); n=3. **(F)** Analysis of cell number, proliferation, and expression of differentiation markers in the experimental con0ditions described in E. Cell number determined with automated cell counter and cell proliferation with CTV staining. Expression of differentiation markers was measured after stimulation with SIINFEKL and brefeldin for 3 hours and is shown as a percentage of live CD8^+^ cells. Histograms are representative flow cytometry plots for each parameter and are pre-gated on live, singlet, CD8^+^ events. The dotted line defines in violin plot graphs the median of 21%-grown cells and in histograms the gate for each marker; n=8-30. **(G)** *In vitro* cytotoxicity assay using OT-I CD8^+^ T cells shown in F. OT-I cells were co-cultured with B16–OVA tumour cells at different effector:target (E:T) ratios and cytotoxicity was assessed after 14-18 hours. **(H)** Cytotoxicity assay obtained according to G performed at 1%O_2_ and shown in dose-response curves (plotted with 95% confidence intervals as shaded areas) determined with non-linear regression ([agonist] vs normalised response). Asterisk represents significantly lower EC_50_ values as compared to 21%; n=4-7. **(I)** Correlation between log_2_ fold change (FC) of cell counts on day 3 (obtained in F) and EC_50_ values (obtained in H) relative to 21% O_2_. All results were pooled from at least two independent experiments and are shown as median ± interquartile range (IQR). Each data point represents an independent animal. * P<0.05, ** P<0.01, *** P<0.001; Holm-Šídák’s multiple comparisons test relative to 21% (E-H) or # P<0.05, ### P<0.001; paired T test (F). Full data and statistical analysis from Figure 1! can be found in Source data 1.

We next wished to determine which of the hypoxia-driven alterations in T cell activation rely on the observed increase in HIF-1α levels elicited by low oxygen tensions (**Figure 1C**). We did this by increasing HIF signalling through pharmacological inhibition of prolyl hydroxylation, utilising the prolyl hydroxylase inhibitor FG-4592 (Chen et al., 2019) (**Figure 1D**). Both low oxygen tensions and FG-4592 act to inhibit proteasomal degradation of HIF-α, leading to HIF-1α accumulation (**Figure 1E**). To facilitate functional characterization, we used CD8^+^ OT-I T cells derived from mice carrying a transgene to express a specific TCR that reacts to the SIINFEKL peptide of ovalbumin (OVA) (Hogquist et al., 1994). Similar to polyclonal T cells, low oxygen conditioning of OT-I cells for 3 days following activation impaired clonal expansion, and increased levels of CD44, CD127, GZMB and IFN-γ (**Figure 1F and S1D**). At the selected concentration (50 μM), FG-4592 increased the expression of the effector molecules GZMB and IFN-γ without impacting the cell expansion, which is relevant for an *ex vivo* protocol of T cell expansion (**Figure 1F**). Importantly, expression of GZMB and IFN-γ was assessed following SIINFEKL rechallenge at day 3, showing that both low oxygen conditioning and FG-4592 could prime T cells with an enhanced capacity to respond to restimulation.

OT-I cells grown in 1% O_2_ showed increased antigen-specific cytotoxicity against B16F10 melanoma tumour cells expressing OVA peptide when compared to wild-type CD8^+^ T cells grown at 21% O_2_ (**Figure 1G-H**). This increased cytotoxicity occurred when assays were conducted at 1% O_2_, 5% O_2_ or 21% O_2_ (**Figure S1E**). Despite belonging to the physiological oxygen range at which cells are exposed to *in vivo*, 5% O_2_ did not impact OT-I cell function under our experimental conditions *in vitro*, nor did treatment with FG-4592 (**Figure 1H**). This might both indicate that only a more pronounced reduction in oxygen levels can functionally modulate T cells, and that the effects of low oxygen conditioning are not fully driven by altered HIF signalling.

Increased cytotoxicity in these populations (illustrated by lower EC_50_ values) was accompanied by decreased cell numbers on day 3 (**Figure 1I**), highlighting that a trade-off relationship exists between CD8^+^ T cell proliferation and effector function when these are induced by low oxygen tensions and increased HIF signalling.

### Modulation of hypoxic response *ex vivo* improves antitumour T cell function *in vivo*

We next compared the effects of hypoxia and HIF signalling modulation on antitumour CD8^+^ T cell function by employing a model of adoptive cell transfer (ACT) (**Figure 2A**). This model allows for an assessment of the long-term effects of temporary exposure of CD8^+^ T cells to low oxygen or PHD inhibition. Wild-type CD45.2^+^ mice were subcutaneously inoculated with OVA-expressing B16 cells, lymphodepleted with cyclophosphamide, and then injected with CD45.1^+^ OT-I CD8^+^ T cells after three days of activation. The use of the CD45.1 and CD45.2 markers allowed us to follow transfused cells after adoptive transfer into experimental recipients (**Figure S2A**).

**Figure 2.**
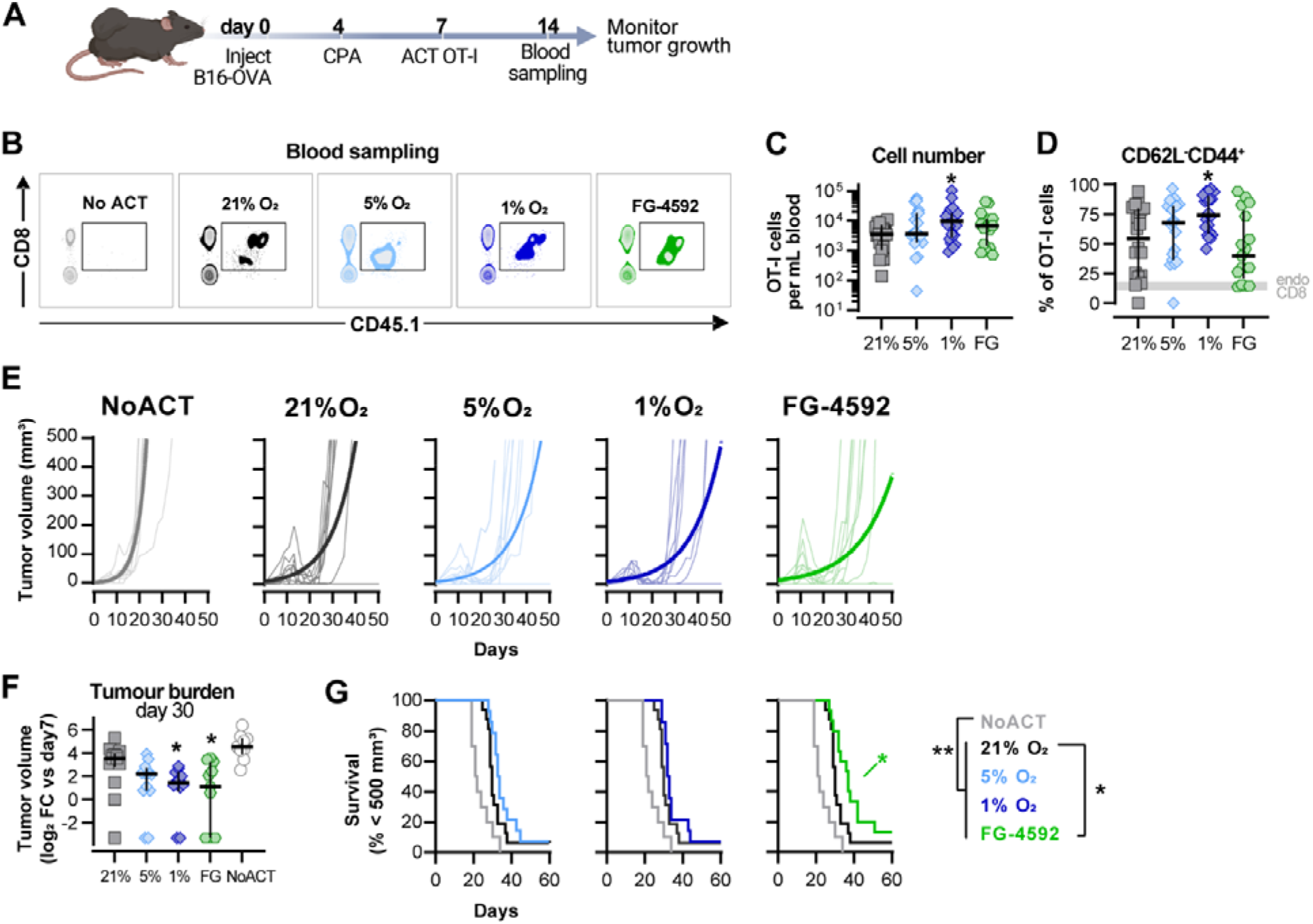
Effects of hypoxia and increased HIF signalling in antitumour function of CD8^+^ T cells. **(A)** Model of adoptive cell transfer (ACT) therapy. CD45.2^+^ C57BL/6j mice were inoculated with B16-OVA cells, lymphodepleted with cyclophosphamide (CPA) and injected with CD45.1^+^ OT-I CD8^+^T cells activated 3 days prior. Peripheral blood was sampled and analysed by flow cytometry. Tumour growth was monitored every 2-3 days until day 60. **(B)** Blood analysis on day 14. Representative flow cytometry plots with events pre-gated on live, singlet, CD45^+^. **(C and D)** Frequency of adoptively transferred OT-I cells per millilitre of peripheral blood (C) and percentage of CD62L^-^CD44^+^ among CD8^+^ CD45.1^+^ in peripheral blood (D) on day 7 after ACT; n=17-20, median ± IQR, each datapoint represents and independent animal. **(E)** Tumour growth curves. Animals not receiving T cells (NoACT) were used as negative controls. Thin lines: individual animals; thick lines: Malthusian growth curve fit; n=7-10. **(F)** Tumour burden on day 30 shown as log_2_ fold change relative to day 7 (day of ACT). The value of 0.1 (corresponding to the smallest log_2_ fold change detected) was added to all conditions to allow the plotting of animals with no tumours; n=9-14, median ± IQR, each datapoint represents and independent animal. **(G)** Survival curves for tumour growth shown in D. Threshold for survival was set to 500mm^3^; n=10-16. All results (except E) were pooled from two independent experiments. * P<0.05, ** P<0.01, *** P<0.001; Kruskal-Wallis test relative to 21% corrected with Dunn’s test (C-D and F) and log-rank (Mantel-Cox) test relative to 21% O_2_ or NoACT groups (G). Full data and statistical analysis from Figure 2 can be found in Source data 2.

Culturing of OT-I CD8^+^ T cells in 1% O_2_ *ex vivo* markedly improved expansion in peripheral blood when the numbers of transferred cells were assayed at 7 days after ACT (**Figure 2B-C**). These results, whereby culture of T cells *ex vivo* in hypoxia gives rise to a striking *in vivo* proliferative advantage, were quite unexpected. Cultivation of T cells *ex vivo* showed that hypoxic stress (i.e., culture in low oxygen) greatly reduces proliferation following activation (**Figure 1B and F**). T cells cultured in 1%O_2_ additionally had a higher proportion of terminally differentiated CD62L^-^CD44^+^ cells 7 days after ACT (**Figure 2D**), following their pattern of increased expression of CD44 and reduced levels of CD62L observed prior to ACT (**Figure S1D**). ACT of OT-I cells cultured in 5% O_2_, or in the presence of FG-4592, did not impact their expansion in the blood or their expression of CD44 or CD62L (**Figure 2B-D**).

Tumour growth was measured for up to 40 days following ACT. The transfer of any OT-I cells delayed tumour growth in comparison with animals not receiving ACT (**Figure 1E-G**). When compared to animals receiving WT OT-I cells grown at 21% O_2_, adoptive transfer of FG-4592-or 1% O_2_-treated OT-I T cells was found to initially decrease tumour burdens (**Figure 2E-F**); while FG-4592-treated cells significantly extended animal survival (**Figure 2G**). Animals receiving OT-I cells conditioned to 5% O_2_ *in vitro* did not show differences relative to control tumour growth (**Figure 2E-G**).

These data show that enhanced effector function of T cells conditioned to low oxygen, or to PHD inhibition *ex vivo* translates to an improved antitumour efficacy *in vivo* following ACT.

### Fine-tuned HIF signalling improves CAR-T cell cytotoxicity against solid tumours

Results shown in Figures 1-2 argue that an increase in HIF signalling through PHD inhibition with FG-4592 can act to improve cytotoxic T cell function in a similar manner to that observed with low oxygen conditioning. To determine whether increase HIF signalling can improve T cell function *in vivo* in CAR therapies and in human T cells, we employed a model of CAR-T cell therapy. We compared the effects of pharmacological inhibition of HIF degradation using FG-4592 *ex vivo*, to the constitutive downregulation of HIF degradation using a vector construct to silence the E3 ubiquitin ligase VHL (**Figure 3A**).

**Figure 3.**
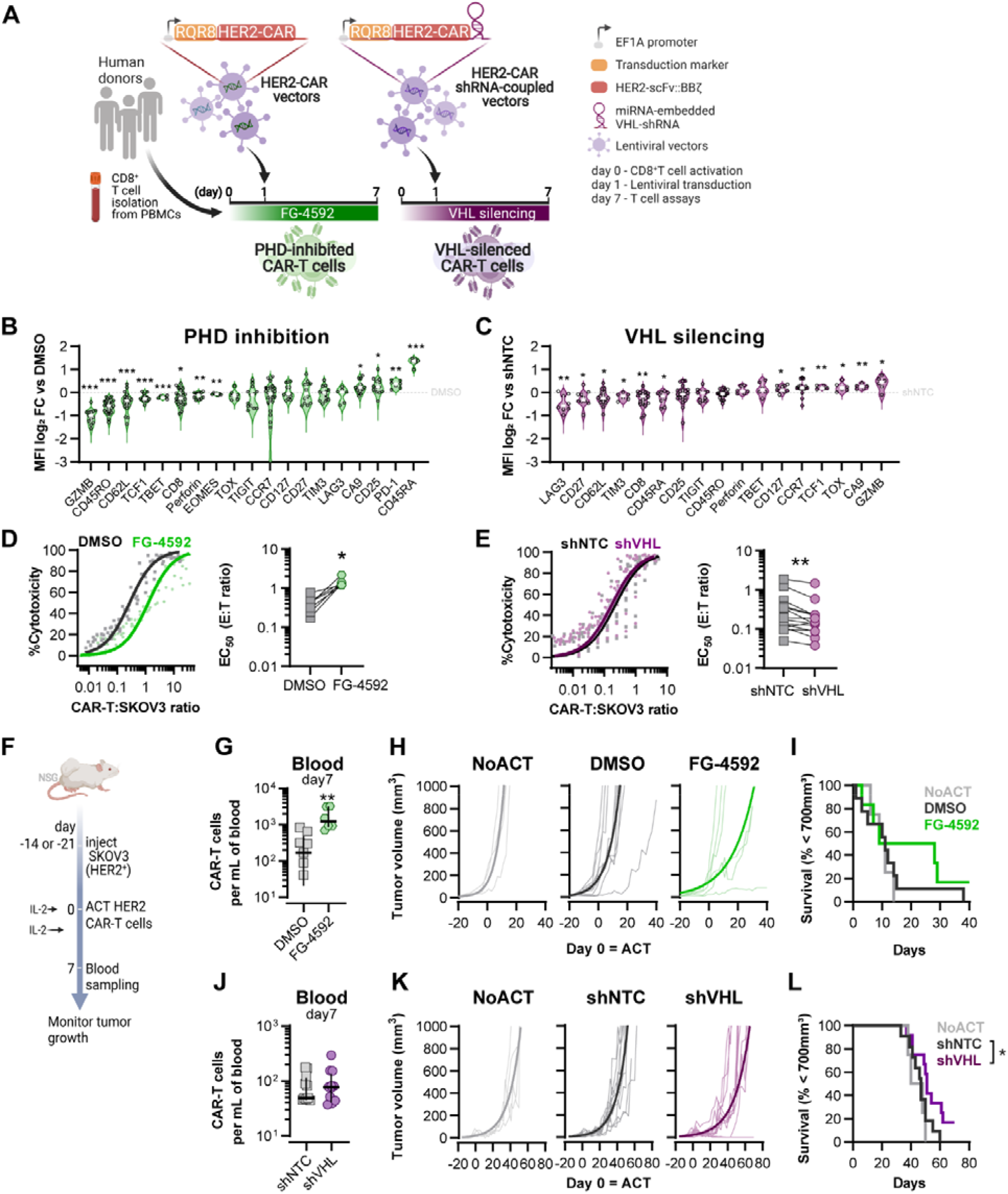
Effects of modified HIF signalling in CART cell cytotoxicity against solid tumours. **(A)** Generation of CAR-T cells. Human CD8^+^ T cells were activated, transduced HER2-CAR vectors the day after and assayed on day 7. To modulate HIF signalling, 50 μM FG-4592 (FG) were added to CAR-T cell cultures or cells were transduced with a HER2-CAR vector containing a miRNA-embedded shRNA against VHL (shVHL). DMSO treatment and transduction with CAR vector containing a non-targeted control sdhRNA (shNTC) were used as controls for FG and shGCDH conditions, respectively. RQR8 was used as transduction marker. **(B and C)** Expression of differentiation markers determined by flow cytometry and shown as log2 fold change in MFI relative to each respective control (horizontal grey line) following PHD inhibition with FG treatment (B) or VHL silencing (D); n=6-30, each data point represents and independent animal. **(D and E)** *In vitro* cytotoxicity assay. DMSO-or FG-treated CAR-T cells (D) and shNTC-or shVHL-expressing CAR-T cells (E) were co-cultured with SKOV3 tumour cells at different effector:target (E:T) ratios and cytotoxicity was assessed after 14-18 hours of co-culture at 1% O_2_. Left: dose-response curves (plotted with 95% confidence intervals represented in shaded areas) determined with non-linear regression ([agonist] vs normalised response); right: EC_50_ values obtained from non-linear regression; n=7-14, each data point represents and independent animal. **(F)** Model of CAR-T cell therapy. NOD.Cg-*Prkdc*^*scid*^*Il2rg*^*tm1Wjl*^/SzJ mice were inoculated with SKOV3 cells and injected with CAR-T cells generated according to A. Peripheral blood was sampled and analysed by flow cytometry. Tumour growth was monitored every 2-3 days until day 40 or 70 for FG and shVHL experiments, respectively. **(G and J)** Frequency of adoptively transferred CAR-T cells per millilitre of peripheral blood on day 7 for DMSO- or FG-treatment (G) and shNTC-or shVHL-silencing (J); n=6-12, median ± IQR. **(H and K)** Tumour growth curves experiment with DMSO-or FG-treated CAR-T cells (H) and shNTC- or shVHL-silenced CAR-T cells (K). Animals not receiving T cells (NoACT) were used as negative controls in both experiments. Thin lines: individual animals; thick lines: Malthusian growth curve fit; n=5-12. **(I and L)** Survival curves for tumour growth shown in H and K. Threshold for survival was set at 700mm^3^. * P<0.05, ** P<0.01, *** P<0.001; One sample T test relative to control (B-C), Wilcoxon matched-pairs signed rank test (D-E), unpaired T test (G) and log-rank (Mantel-Cox) test relative to shNTC (L). Full data and statistical analysis from Figure 3 can be found in Source data 3.

For this we used HER-2 CAR-T cells targeted to a solid tumour model of SKOV3 human ovarian epithelial cancer. RQR8 was used as a transduction marker after confirming co-expression with CAR molecules at the surface (**Figure S3A**). To achieve VHL silencing, we engineered a CAR vector which is coupled to the expression of a microRNA embedded Vhl shRNA (**Figure 3A**). For expression of shRNAs, the commonly used RNA polymerase III promoters (e.g., U6), which can drive expression of small RNAs, often result in abundant production of small RNAs with heterogeneous 5’ ends; these can cause a deleterious increase of off-target effects. High levels of small RNA molecules can also become toxic to cells by saturating factors required for generating endogenous non-coding RNAs. To avoid these issues, shRNA-hairpins can be flanked with an optimized sequence of miR-30 (Fellmann et al., 2013) and can be expressed under the control of an RNA II polymerase promoter (e.g., EF1α). This promoter enables the coexpression of protein and shRNA, which are processed like endogenous miRNAs. In a proof of concept experiment, CD5 (highly expressed in lymphocytes) was successfully downregulated in T cells transduced with CD5-shRNA vectors, as confirmed by flow cytometry in RQR8+ cells, when compared to RQR8-events (non-transduced) or with cells transduced with non-targeted control (NTC) shRNAs (**Figure S3A-B**). This occurred whether the vector co-expressed a HER2-CAR sequence along with the shRNA or not. Silencing of CD5 expression was stable until at least day 12 after transduction (**Figure S3C**).

Transduction of human CD8^+^ T cells with VHL-targeted shRNA (shVHL) using an engineered CAR-vector reduced VHL expression by approximately 40% (**Figure S3D**); this was sufficient to detectably increase HIF levels (**Figure S3E**). Culturing T cells for seven days with PHD inhibition, or silencing VHL by shRNA, both increased expression of the HIF target gene Carbonic Anhydrase 9 (Ca9), and reduced expression of CD62L relative to non-treated controls (**Figure 3B-C**). However, expression of most markers of differentiation was remarkably different between these two cell populations, and relative to controls. In particular, GZMB, TOX and TCF1 were decreased by pharmacological PHD inhibition, but were increased in the T cells transduced with the VHL silencing vector. *In vitro*, while FG-4592 treatment impaired cytotoxicity of CAR-T cells, shVHL-expressing CAR-T cells showed enhanced tumour cell killing capacity (**Figure 3D-E and S4F**).

We next employed a model of ACT using these HER2-CAR-T transduced human T cells. This allowed for an assessment of long-term effects of both temporary and constitutive modulation of the HIF pathway achieved through the pharmacological FG-4592 treatment *ex vivo;* or via the vector-induced silencing of VHL. Immunocompromised (NSG) mice were subcutaneously inoculated with HER2-expressing SKOV3 human cancer cells and then injected with RQR8^+^ HER2-CAR expressing human donor CD8^+^ T cells, generated as shown in Figure 3A. After ACT was delivered to recipient animals, FG-treated CAR-T cells expanded to a greater extent in the blood relative to DMSO-treated control T cells (**Figure 3G**). Tumour growth analyses showed that, relative to animals not receiving CAR-T cells, FG-4592 treated CAR-T cells delayed tumour growth in 50% of the animals compared to 9% of the animals in the control T cell group (**Figure 3H**). However, the FG-4592 treatment group did not statistically significantly improve animal survival, in spite of having one animal with a complete cure (**Figure 3I**). VHL silencing by the shRNA vector did not impact cell expansion *in vivo* (**Figure 3J**). However, VHL-silenced HER2-CAR-T cells were able to decrease tumour growth significantly, and an improved animal survival relative to NTC-expressing CAR-T cells was seen, clearing tumours in 2 of the 11 animals analysed (**Figure 3K and L**).

These data show that an increase in HIF signalling can potentiate T cell responses *in vivo*, as seen both by increased expansion of FG-treated cells, and through increased tumour killing capacity of VHL-silenced cells; indicating HIF can therapeutically increase human CAR-T cell function.

### Short hypoxic conditioning is sufficient to shape T cell differentiation

We found that low oxygen conditioning increases HIF stabilisation in activated CD8^+^ T cells (**Figure 1C**). After establishing that protocols of HIF stabilisation in T cells with PHD inhibition or VHL silencing can modulate differentiation and function, we became interested in understanding whether the duration of exposure to low oxygen would also have functional consequences for CD8^+^ T cells. For this, we conditioned human CD8^+^ T cells for 1, 3, or 7 days at 1% O_2_ during a 7-day culture protocol and compared their differentiation to that of cells cultured exclusively under ambient (21% O_2_) conditions (CT) (**Figure 4A**).

**Figure 4.**
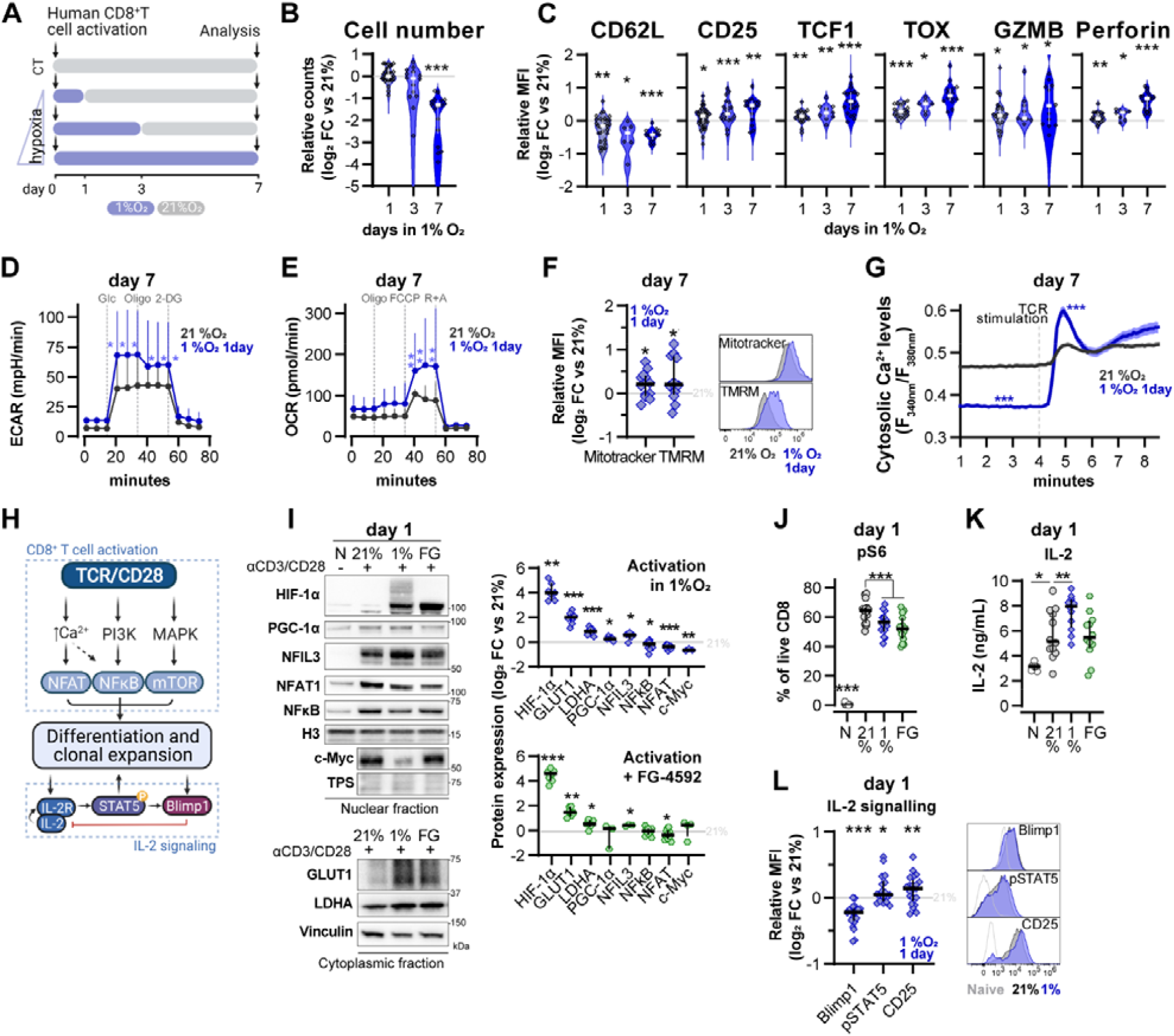
Impact of increasing exposure to hypoxia in human CD8^+^ T cell activation and differentiation. (**A**) Human CD8^+^ T cells were activated in 1 %O_2_ for 1, 3 or 7 days. Cells continuously grown in 21% were used as control (CT). (**B**) Number of CD8^+^ T cells after 7 days of increasing time of exposure to 1%O_2_ as shown in A. Cell counts assessed by flow cytometry using counting beads and presented as log_2_ FC relative to 21% cultures; n=14-29. **(C)** Expression of differentiation markers shown as log2 FC in MFI relative to 21% CT (horizontal grey line) following conditioning to 1% O_2_ as described in A; n=8-38. **(D and E)** Seahorse analysis of activated human CD8^+^ T cells exposed for 1 day to 1%O_2_ as shown in A. ECAR determined after injection of glucose (Glc), Oligomycin (O) and 2-DG (D). OCR determined after injection of Oligomycin (O), FCCP (F) and rotenone+antimycin A (R+A) (E); n=12. **(F)** Signal of mitotracker DeepRed and TMRM in 7 day-cultured cells exposed to 1%O_2_ for 1 day as shown in A. Left: log_2_ FC relative to 21% controls; Right: histograms representative for cells activated in 21% O_2_ or in 1% O_2_ for 1 day (blue). **(G)** OT-I T cells were activated for 24 hours in 1%O_2_ with SIINFEKL and expanded for 6 days in 21% O_2_ before single-cell cytosolic Ca^2+^ was measured by microscopy using the radiometric dye Fura-2AM (F_340nm_/F_380nm_) in presence of 1 mM Ca^2+^ at day 7. Single-cell time-lapse imaging was performed at 5 s intervals with baseline fluorescence measured for 4 min, followed by another 6 min reading after addition of SIINFEKL. Number of cells analysed (21% grey, n=249; 1%1day blue, n=128) are derived from multiple coverslips and 2 OT-I donor spleens; mean ± SEM. **(H)** TCR signalling. **(I)** Protein analysis 24h after activation of human CD8^+^ T cells with anti-CD3/CD28 dynabeads in 21% O_2_, 1% O_2_ or 21% O_2_ + 50μM FG-4592 (FG). Non-activated cells (N) were used as negative control. Histone 3, Lamin B, HDAC or total protein stain (TPS) were used as loading control for nuclear fraction proteins and vinculin was used as loading control for the cytoplasmic fraction. Left: representative immunoblots; right: log_2_ FC in protein expression of cells activated in 1% (top) or with FG (bottom) relative to 21% controls; n=3-12. **(J)** Flow cytometry analysis of the proportion of cells positive for phospho-S6 Ribosomal Protein (Ser235/236, pS6) for the same experimental conditions described in H; n=10-15. **(K)** IL-2 ELISA performed on supernatant of cells activated for 1 day as described in I; n=6-12. **(L)** Flow cytometry analysis of Blimp1, phopho STAT5 (Y694, pSTAT5) and CD25 in cells activated in 1% O_2_ for 24 hours. Left: MFI log_2_ fold change relative to 21% O_2_. Right: representative histograms for naive (light grey) and cells activated in 21% (dark grey) and 1% O_2_ (dark blue). All results were pooled from at least two independent experiments and are shown as median ± IQR (except G). Apart from D and E, each data point represents an independent donor; * P<0.05, ** P<0.01, *** P<0.001; Wilcoxon matched-pairs signed rank test (B-C and L), Šídák’s multiple comparisons test relative to 21% (D-E, J-K) and unpaired (G) and paired (F and I) T tests. Full data and statistical analysis from Figure 4 can be found in Source data 4.

As was seen in murine CD8^+^ T cells, incubation in low oxygen conditions decreases human CD8^+^ T cell expansion (**Figure S4A**). Longer periods spent at 1% O_2_ correlated with the strongest reduction in cell numbers when compared to 21% O_2_-grown cells (**Figure 4B**). However, the expression of differentiation markers elicited by 1, 3, and 7 days of 1% O_2_ conditioning was remarkably similar, although the magnitude of differential expression increased with longer exposure to 1% O_2_ (**Figure 4C and S4B**). A similar trend was found when we carried out exposures to 5% O_2_ conditioning (**Figure S4C-D**). On the other hand, incubation in low oxygen conditions (1% or 5% O_2_) during the last 3 days of culture had little effect (**Figure S4E-F**), despite leading to the highest levels of HIF-1α on day 7 (**Figure S4G**). This indicates that oxygen levels at the time of activation are those that can profoundly shape T cell differentiation.

In fact, a single day of 1% O_2_ conditioning, occurring at the time of activation, followed by 6 days of 21% O_2_ culture, was sufficient to elicit a classical hypoxia/HIF-driven metabolic adaptation in T cells (Papandreou et al., 2006) as characterized by an increased glycolytic rate (**Figure 4D**). This short-term exposure to hypoxia at the time of activation led to increased oxygen consumption rates at baseline and at maximal capacity after FCCP addition (**Figure 4E**). The change in oxygen consumption rates was accompanied by both an increased mitochondrial DNA copy number (**Figure S4H**) and by increased mitotracker (indicating increased mitochondrial mass) and increased TMRM staining (indicating a shift in mitochondrial membrane potential) (**Figure 4F**) when compared to cells continuously grown in 21% O_2_, as measured at 7 days following initial activation. Single-cell fluorescent microscopy using the ratiometric calcium dye Fura-2AM shows that conditioning to 1% O_2_ exclusively during the first 24 hours of activation of OT-I cells results in lower baseline intracellular calcium levels when compared to cells continuously cultured in ambient oxygen. However, restimulation with SIINFEKL at day 7 induced a much higher store-operated calcium entry (SOCE) in the cells exposed to 1% O_2_ (**Figure 4G and S4I**). Activation of SOCE and the concomitant rise in cytosolic calcium is a key event in TCR signalling, leading to activation of calcineurin and translocation of Nuclear Factor of Activated T-cells (NFAT) to the nucleus (Serfling et al., 1995); indeed, we do see an accompanying increase in NFAT nuclear translocation in hypoxia conditioned group at day 7 (**Figure S4J**).

To understand the impact of these transient low oxygen tensions on T cell activation, we next characterized the expression of transcription factors and other proteins related to TCR signalling and IL-2 signalling in CD8^+^ T cells after 24 hours of activation in 1% O_2_ (**Figure 4H**). We compared the alterations elicited by 1% O_2_ to those seen during differentiation in ambient oxygen (21%) cultured T cells, and to FG-4592-treated T cells (**Figure 4I and S4K**). Nuclear translocation of NFAT was reduced by 1% O_2_ and PHD inhibition, whereas the levels of Nuclear factor kappa B (NFkB) were only reduced in the nuclear fraction of 1% O_2_ activated T cells. Both NFAT1 and NFkB are crucial for T cell differentiation (Serfling et al., 1995). Levels of peroxisome proliferator-activated receptor gamma coactivator 1-alpha (PGC-1α, a master regulator of mitochondrial biogenesis (Liang and Ward, 2006)) and c-Myc (a primary mediator of glycolysis in T cells (Wang et al., 2011)) were respectively increased and decreased by activation during a short period of in 1% O_2_ (**Figure 4E-G**).

The increase in HIF-1α levels following activation was augmented by exposure to 1% O_2_ or PHD inhibition (**Figure 4I**). This correlated with increased levels of the known HIF targets glucose transporter 1 (GLUT1) and lactate dehydrogenase A (LDHA), as well as the Nuclear Factor Interleukin 3 Regulated (NFIL3), previously reported to be increased following ectopic expression of HIF-α or culture in 1% O_2_ for 24 hours (Ross et al., 2021; Veliça et al., 2021). As NFIL3 has been linked to perforin production by NK cells, this transcription factor could underpin the increased T cell production of effector molecules following low oxygen conditioning or increased HIF signalling (Rollings et al., 2018). Moreover, 1% O_2_ culture, and PHD inhibition both decreased mTOR activation, as assessed by flow cytometry analysis of the mTOR target Phospho-S6 Ribosomal Protein (Ser235/236)S6 (**Figure 4J**). mTOR signalling is important for the growth and survival of activated T cells, and is also involved in the metabolic rewiring of T cells by driving the expression of c-Myc and HIF-1α (Finlay et al., 2012; Wang et al., 2011). Hypoxia and HIF have been previously shown to dampen mTOR signalling (Wouters and Koritzinsky, 2008), which could explain the above-mentioned reduced expression of c-Myc in T cells activated under low oxygen tensions.

T cell activation in low oxygen increased the production of IL-2 when compared to cells activated in ambient oxygen (**Figure 4K**). While FG-4592 treatment did not impact IL-2 production in T cells, we have previously found that *Il2* gene expression is increased by ectopic HIF-1α in mouse T cells (Veliça et al., 2021). In fact, we observed that the increase of IL-2 production following 24 hours of T cell activation in hypoxia was accompanied by an increase in STAT5 phosphorylation (Y694) and in the expression of IL-2 receptor subunit CD25 (**Figure 4L**). STAT5 is a positive regulator of IL-2 signalling previously shown to be induced by hypoxia in non-immune cells (Joung et al., 2003), whereas CD25 is one of the most consistently upregulated targets in T cells cultured under low oxygen tensions or following increased HIF signalling (Figure 1A, 3C and 5C). CD25 upregulation by hypoxia in T cells has also been shown by others (Caldwell et al., 2001; Ross et al., 2021). In contrast with pSTAT5 and CD25, expression of Blimp-1 (a transcriptional repressor of IL-2 signalling (Malek, 2008) and PGC-1α (Malek, 2008; Scharping et al., 2021) were decreased in CD8^+^ T cells following 24 hours of activation in 1%O_2_ (**Figure 4L**). This could both explain the increase in IL-2 production and the decreased PGC-1α expression obtained following activation in low oxygen. Blimp1, a known HIF target (Chiou et al., 2017), is increased by ectopic expression of HIFα in mouse T cells (Veliça et al., 2021). However, its expression is increased by T cell activation and TCR signalling (Martins and Calame, 2008), which is dampened by a 24h activation in 1% O_2_ (as seen by decreased nuclear translocation of NFAT and NfKB) and could underlie the observed decrease in Blimp1 expression.

Together, these findings suggest that low oxygen levels at the time of activation specifically shape T cell differentiation. Most, but not all, alterations elicited on T cell activation signalling by low oxygen conditioning can be recapitulated by PHD inhibition, indicating that HIF drives hypoxia-related reprogramming of T cells at the time of T cell activation.

### Short hypoxic conditioning improves T cell function against solid tumours

Lastly, we assessed the effect of the duration of low oxygen stimulation on CAR-T cell function. For this, we conditioned HER2- and CD19-CAR-T cells with low oxygen tensions for 1 day (**Figure 5A**) or 3 days (**Figure S5A**) during a 7-day culture protocol. Cytotoxic CAR-T cell function was not improved by the more prolonged 3-day exposure to 5 or 1% O_2_ (**Figure S5B-E**). However, a single day of 1% O_2_ conditioning followed by 6 days of culture in 21% (1% O_2_ conditioning for 1 day) enhanced the killing capacity of both HER2- and CD19-CAR-T cells, with a significantly increased secretion of IFN-γ (**Figure 5B-C**). Conditioning CAR-T cells for 1 day at 5% O_2_ did not impact cytotoxic function (**Figure S5F-G**).

**Figure 5.**
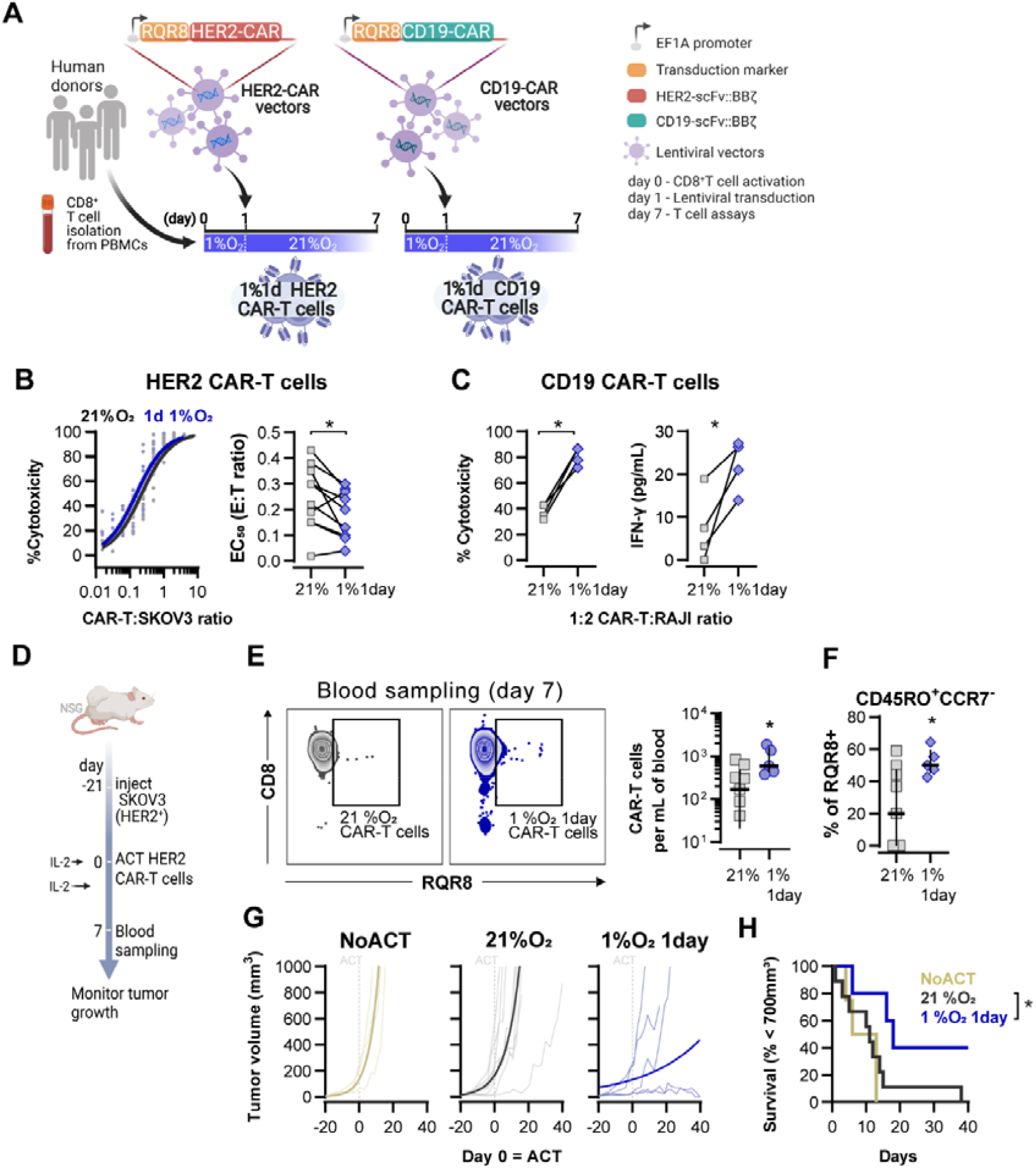
Effects of short hypoxic conditioning in CAR-T cell function against solid tumours. **(A)** Generation of CAR-T cells. Human CD8^+^ T cells were activated, transduced with HER2- or CD19-CAR vectors the day after and assayed on day 7. Cells were activated in 1% O_2_ for 24 hours and swapped to 21% until day 7. CAR-T cells continuously maintained in 21% O_2_ were used as controls and RQR8 was used as transduction marker. **(B)** *In vitro* cytotoxicity assay with HER-2 CAR-T cells cultured according to A. CAR-T cells were co-cultured with SKOV3 tumour cells at different effector:target (E:T) ratios and cytotoxicity was assessed after 14-18 hours of co-culture at 1% O_2_. Left: dose-response curves (plotted with 95% confidence intervals represented in shaded areas) determined with non-linear regression ([agonist] vs normalised response); right: EC_50_ values obtained from non-linear regression; n=6. **(C)** *In vitro* cytotoxicity assay with CD19 CAR-T cells cultured according to A. CAR-T cells were co-cultured with RAJI tumour cells at a E:T ratio of 2:1 and cytotoxicity was assessed after 14-18 hours of co-culture at 5% O_2_. Left: cytotoxicity relative to cells transduced with vector control containing RQR8 alone; right: IFN-γ ELISA using supernatants of the co-culture between CAR-T cells and RAJI; n=4. **(D)** Model of CAR-T cell therapy. NOD.Cg-*Prkdc*^*scid*^*Il2rg*^*tm1Wjl*^/SzJ mice were inoculated with SKOV3 cells and injected with HER-2 CAR-T cells generated according to A. Peripheral blood was sampled and analysed by flow cytometry. Tumour growth was monitored every 2-3 days until day 40. **(E)** Blood analysis on day 14. Left: representative flow cytometry plots with events pre-gated on live, singlet, human CD45^+^. Right: frequency of adoptively transferred CAR-T cells per millilitre of peripheral blood; n=5-7. **(F)** Percentage CD45RO^+^CCR7^-^ cells among RQR8+ cells in peripheral blood on day 7 after ACT; n=5-7. **(G)** Tumour growth curves. Animals not receiving T cells (NoACT) were used as negative controls. Thin lines: individual animals; thick lines: Malthusian growth curve fit; n=5-12. **(H)** Survival curves for tumour growth shown in F. Threshold for survival was set at 700mm^3^. * P<0.05, ** P<0.01, *** P<0.001; Wilcoxon matched-pairs signed rank test relative to control (B), Mann-Whitney test relative to control (C and E-F) and log-rank (Mantel-Cox) test relative to 21% (H). Full data and statistical analysis from Figure 5 can be found in Source data 5.

Adoptive transfer of HER2-CAR-T cells to NSG mice bearing HER2^+^-SKOV3 tumours was used to functionally compare CAR-T cells exposed for 1 day to 1%O_2_, to cells continuously cultured in 21% O_2_ for 7 days prior to ACT (**Figure 5D**). We found that a short 1% O_2_ conditioning *ex vivo* during activation improved CAR-T expansion *in vivo*, as assessed by analysis of the blood from recipient animals 7 days after ACT (**Figure 5E**). Additionally, 1% oxygen for 1 day conditioned CAR-T cells were found to have a higher proportion of effector memory cells (CD45RO^+^CCR7^-^) in peripheral blood when compared to 21% oxygen conditioned CAR-T cells (**Figure 5F**). A single day of 1% O_2_ conditioning of HER2-CAR-T cells significantly enhanced their immunotherapeutic function against SKOV3 tumours, as shown by significantly decreased tumour growth rates, and by increased animal survival, relative to animals receiving 21% O_2_ treated CAR-T cells (**Figure 5G-H**).

These data indicate that oxygen tensions during activation can act to permanently shift T cell differentiation and function, with a short period of low oxygen conditioning allowing a shift in T cell differentiation, and an increase in cytotoxic capacity, both *ex vivo* and *in vivo*.

## Discussion

Research investigating the role of hypoxia in CD8^+^ T cell-mediated immunity has generally focused on the immunosuppressive consequences of oxygen depletion in the tumour microenvironment (Chouaib et al., 2017; Jayaprakash et al., 2021). While tumour hypoxia correlates with poor patient survival and reduced T cell infiltration(Bertout et al., 2008; Hatfield et al., 2015), data presented here show that the effects of low oxygen tensions on CD8^+^ T cells are not solely inhibitory, and can significantly augment aspects of adaptive immune responses.

In T cells, cognate antigen stimulation of the T cell receptor (TCR) triggers an acute HIF-1α upregulation (even at the high oxygen tensions present in tissue culture), which drives a transcriptional program that supports both effector differentiation and aerobic glycolysis (Finlay et al., 2012; Nakamura et al., 2005; Palazon et al., 2017). In CD8^+^ T cells, HIF-1⍰ and HIF-1ɑ (but not HIF-2ɑ) deletion resulted in incomplete effector differentiation and reduced expression of glycolytic genes((Clever et al., 2016; Doedens et al., 2013; Finlay et al., 2012; Liikanen et al., 2021; Palazon et al., 2017; Veliça et al., 2021)). Deletion of the various negative HIF regulators, the HIF inhibitor (FIH/HIF1AN) or the von Hippel-Lindau disease tumour suppressor (VHL) protein, or the prolyl hydroxylase domain-containing proteins (PHD); or overexpression of modified, negative regulator-insensitive HIF-ɑ proteins, exacerbates the terminal effector program (Clever et al., 2016; Doedens et al., 2013; Finlay et al., 2012; Liikanen et al., 2021; Palazon et al., 2017; Veliça et al., 2021). In adoptive cell transfer experiments, there are clearly complexities found in the role of HIF in overall T cell function during overexpression: HIF-1ɑ deletion and overexpression of VHL- and FIH-insensitive HIF-ɑ impaired CD8^+^ T cell antitumour function, while VHL deletion and overexpression of VHL-insensitive and FIH-sensitive HIF-2ɑ improved cytotoxicity (Liikanen et al., 2021; Palazon et al., 2017; Veliça et al., 2021).

CD8^+^ T cell activation mediated an increase in HIF signalling that we found to be amplified by lower oxygen tensions. *Ex vivo* low oxygen conditioning of CD8^+^ T cells caused an increase in the production of effector molecules and cytotoxic function, but also severely impaired cell proliferation. This coupling of increased HIF, increased cytotoxic function, and decreased cell growth is a finding which is consistent with previous reports (Gropper et al., 2017; Ross et al., 2021; Xu et al., 2016). Having previously shown that HIF-α deletion impairs CD8^+^ T cell function (Palazon et al., 2017), here we describe that HIF-α accumulation, achieved through pharmacological PHD inhibition via FG-4592 or silencing of VHL, can boost antitumour CD8^+^ T cell function. Pharmacological inhibition of PHDs *ex vivo* has been shown to improve function in DMOG-treated CD4^+^ T cells (Clever et al., 2016). Here we show that similar to conditioning CD8^+^ T cells to low oxygen tensions before ACT, a transient boost in HIF signalling through *ex vivo* treatment with the PHD inhibitor FG-4592 improved the expansion and cytotoxicity of adoptively transferred tumour-specific mouse and human CD8^+^ T cells. This argues that oxygen tensions and/or HIF levels during T cell priming can have long-lasting functional effects on CD8^+^T cells.

Perhaps most surprisingly, we show here that while long periods of low oxygen conditioning strongly reduced T cell expansion, the observed oxygen-dependent increases in effector function occurred regardless of whether the cells were cultured for short or long periods of hypoxia, as long as the hypoxic exposure was coincident with activation. This is best illustrated by the fact that a single day of low oxygen conditioning during T cell activation, followed by expansion in higher oxygen tensions, was sufficient to significantly alter T cell differentiation and metabolism and to improve the antitumour efficacy of CAR-T cells *in vivo*.

24 hours of TCR stimulation and co-stimulation in 1% O_2_ profoundly altered T cell activation. This short period of hypoxia at the time of activation reduced nuclear levels of NFAT1, NFkB and c-Myc; decreased mTOR signalling; increased expression of HIF-1α, GLUT1, LDHA, NFIL3 and PGC-1α; upregulated IL-2 signalling by increasing IL-2 production, CD25 expression and STAT5 phosphorylation while reducing Blimp1 protein levels. After 6 days of ambient oxygen culture, T cells that had been conditioned to low oxygen for the first 24 hours of activation showed increased mitochondria biomass, enhanced maximal glycolytic and respiratory rates, and heightened SOCE following TCR restimulation. Clearly these data demonstrate a previously unappreciated role for oxygen in the process of activation and in persistently altering differentiation of CD8^+^ T cells.

The microenvironment, and the amount of available oxygen, at the time of antigen recognition thus has a highly persistent and determinative effect on T cell function, and this likely represents a novel and important aspect of T cell programming that needs to be better understood. Many of these changes are HIF-driven, but some may well be related to other immunometabolic determinants present in the microenvironment of antigen recognition and activation. The potential for utilizing these findings for short and easily achievable alterations in how therapeutic T cells are cultured presents an important new opportunity for improving efficacy in immunotherapy.

## Materials and methods

### Animals

C57BL/6J (CD45.2) animals were purchased from Janvier Labs. Donor TCR-transgenic OT-I mice (JAX #003831, (Hogquist et al., 1994) were crossed with mice bearing the CD45.1 congenic marker (JAX #002014, (Janowska-Wieczorek et al., 2001). All experiments were performed with age and sex-matched littermate controls. All animal experiments were approved by the regional animal ethics Committee of Northern Stockholm, Sweden.

### Cell lines

B16-F10 was originally purchased from ATCC (CRL-6475) and genetically modified to express ovalbumin, eGFP and neomycin phosphotransferase (Veliça et al., 2021). The resulting ovalbumin expression B16F10 cells were cultured in DMEM high glucose with pyruvate (11995065 ThermoFisher) containing 0.75 mg/mL G418 sulfate (10131027, ThermoFisher). HEK293 was a gift from Prof. Dantuma (Karolinska Institute, Stockholm) and cultured in DMEM high glucose with pyruvate. SKOV3 was purchased from ATCC (HTB-77) and cultured in McCoy’s 5A Medium (16600082, ThermoFisher). Raji-GFP-Luc were purchased from Biocytogen (B-HCL-010) and cultured in complete RPMI (21875, Thermo Fisher). All media was supplemented with 1% penicillin streptomycin (15140122 ThermoFisher) and 10% fetal bovine serum (FBS, A3160802 ThermoFisher). Cell lines were frozen at low passage number (<5) in DMEM containing 10% DMSO and were typically passaged 3 to 4 times between thawing and experimental use.

### T-cell isolation and activation and restimulation

Splenic murine CD8^+^ T lymphocytes were purified with either positive and negative CD8 microbeads (130-117-044 and 130-104-075 Miltenyi, respectively) by magnetic-activated cell sorting (Miltenyi). Activation was done in complete RPMI supplemented with 55 μM 2-ME (21985023, Gibco) and anti-mouse CD3/CD28 dynabeads (11453D, ThermoFisher) at a 1:1 cell-to-bead ratio. Purified OT-I CD8^+^ T cells were activated with 0.1-1 μg/mL of the OVA-derived peptide SIINFEKL (ProImmune) or with anti-mouse CD3/CD28 dynabeads. CD8^+^ T cells were expanded in the presence of 10 U/mL recombinant human IL-2 (11147528001, Sigma). Human CD8^+^ T cells were purified from donor PBMCs (NHSBT or Karolinska Hospital) by positive CD8 magnetic bead cell sorting (130-045-201, Miltenyi) and activated in complete RPMI supplemented with 30 U/mL IL-2 with anti-human CD3/CD28 dynabeads (11131D, ThermoFisher) at a 1:1 cell-to-bead ratio. T cells were pre-incubated for at least 2 hours in the different oxygen tensions of with FG-4592 before activation and were cultured at a density of approximately 5-10 x10^5^ cells per ml per cm^2^.

### Flow cytometry

Single cell suspensions were stained with Near-IR Dead Cell Stain Kit (Thermo Fisher) followed by surface and intracellular staining with fluorochrome-labelled antibodies (Table S1). Staining of cytoplasmic and nuclear antigens was performed using the Fixation/Permeabilization kit (BD Biosciences) and the Transcription Factor buffer set (BD Biosciences), respectively. To measure IFN-γ and TNF-α production, mouse OT-I T cells were incubated with RPMI supplemented with 1 μg/mL SIINFEKL with and treated with 5 μg/mL brefeldin 4 hours before intracellular staining and flow cytometry analysis. For proliferation assays, cells were loaded with CellTrace Violet (Thermo Fisher) according to manufacturer’s instructions. For mitochondria analysis, cells were loaded with 25nM Mitotracker DeepRed (invitrogen) or 30nM TMRM (invitrogen) for 20 min at 37oC. Samples were acquired in FACSCanto II (BD Biosciences) or in Aurora (Cytek Biosciences) flow cytometers and data analysed with FlowJo version 10. Transduced cells were sorted on an Aria III (BD Biosciences) following the surface antigen staining described above.

### Western blotting

Total cell pellets were lysed with urea-tris buffer (8 M urea, 50 mM Tris-HCl (pH=7.5), 150 mM β-mercaptoethanol), sonicated twice for 45 sec intercalated with 1 min incubation on ice and centrifuged at 14000 x g, 4 ºC for 15 min. Histones were extracted with Histone Extraction Kit (ab113476, Abcam). Nuclear and cytoplasmic extracts from CD8^+^ T cells were obtained with the NE-PER Nuclear and Cytoplasmic Extraction Reagents (78833, ThermoFisher). The Revert 700 Total Protein Stain (LICOR) was used for analysis of total protein stain. Proteins (15-30 μg) were separated by SDS-PAGE and transferred to PVDF membranes before being probed with primary antibodies at a 1:1000 dilution (Table S2). Following manufacturer’s instructions, protein signal was detected using infra-red labelled secondary antibodies in an Odyssey imaging system (LI-COR) or using horseradish peroxidase-conjugated secondary antibody (R&D systems) and ECL Prime (GE Healthcare) (imaged with an iBrightCL1000 (ThermoFisher).

### In vitro cytotoxicity assay

10000 B16F10-OVA or SKOV3 cells were seeded per well in 96 well plates (flat bottom, Costar) and co-cultured for a minimum of 14 hours with varying ratios of mouse CD8^+^ OT-I or human CD8^+^RQR8^+^ HER2-CAR-T cells, respectively. Wells were washed twice with PBS to remove T cells and the number of remaining target cells was determined by culturing with 10 μg/mL resazurin (Sigma) for 2 hours and measuring fluorescence signal in a plate reader. Cytotoxicity was calculated relative to wells with no T cells added (positive control) and to wells with no cancer cells added (negative control). 10000 Raji cells were co-cultured with varying ratios of CD8^+^RQR8^+^ CD19-CAR-T cells for a minimum of 14 hours and cytotoxicity was assessed by flow cytometry. The ratio of Raji cells to CountBright Absolute counting beads (Thermo Fisher) was used to calculate cytotoxicity. To determine specific cytotoxicity, data was normalised to the cytotoxicity of VC-transduced CD8^+^ T cells of the respective donor.

### Adoptive cell transfer experiments

8 to 15-weeks old female C57BL/6j CD45.2^+^ mice were inoculated subcutaneously with 5×10^5^ B16-F10-OVA and conditioned 4 days later with peritoneal injection of 300 mg/kg cyclophosphamide (Sigma). On day 7, 0.5-1×10^6^ CD45.1^+^ OT-I CD8^+^ T cells activated for 3 days were peritoneally injected. 8 to 15-weeks old female NOD.Cg-*Prkdc*^scid^*Il2rg*^tm1Wjl^/SzJ mice were inoculated subcutaneously with 1×10^6^ SKOV3 and injected 21 to 7 days later with 5x10^5^ RQR8^+^ human CD8^+^ CAR-T cells expanded for 7 days *ex vivo*. 100U of IL-2 was peritoneally injected in NSG mice on the day of ACT and 3 days later. Animals were assigned randomly to each experimental group and tumour measurements were blinded. Tumour volume (a×b×b/2 where a is the length and b is the width) was measured every 2-3 days with electronic callipers until experimental end date as specified in figure legend.

### Vectors

DNA encoding a codon-optimized polycistronic peptide composed of RQR8 and anti-human HER-2 (clone 4D5) or anti-human CD19 (clone FMC63) interspersed with picornavirus T2A and furin cleavage sequences was synthesized by GeneScript. RQR8, used as the transduction marker, is a chimeric surface protein composed with domains from CD34 (for detection and purification with clone QBEND/10), CD8 (for anchoring at the cell surface) and CD20 (for depletion *in vivo* with anti-CD20 mAb rituximab) (Philip et al., 2014).

MicroRNA embedded shRNAs were generated as previously described (Ros and Gu, 2016). Briefly, 97-mer oligonucleotides (IDT Ultramers) coding for the respective shRNAs (Moffat et al., 2006) were PCR amplified using 10uM of the primers miRE-XhoI-fw (5′-TGAACTCGAGAAGGTATATTG CTGTTGACAGTGAGCG-3′) and miRE-EcoRI-rev (5′-TCTCGAATTCTAGCCCCTTGAAGT CCGAGGCAGTAGGC-3′), 0.5 ng oligonucleotide template, and the Q5 High-Fidelity 2X Master Mix (NEB), and cloned HER-2 CAR vectors containing the miRE scaffold sequence. All coding sequences were cloned into pCDCAR1 (Creative-Biolabs). Third generation lentiviral transfer helper plasmids were obtained from Biocytogen. All sequences are available in Table S3.

### Lentiviral transductions

For generation of lentiviral particles, 5×10^6^ HEK293 cells were plated in 15 cm petri dishes and transfected the day after with 50 μL FuGENE (E2311, Promega), 10 μg CAR-encoding vectors and 3.3 μg of each 3rd generation lentivirus helper vectors (CART-027CL, Creative-Biolabs). Supernatant media containing lentiviral particles was harvested 48 hours after transfection and used fresh or stored at -80 °C. Lentiviral supernatants were spun onto non-treated 24-well plates, coated with 30 μg/mL Retronectin reused up to three times (T100B, Takara), at 2000 x g for 2 hours at 32°C and replaced with activated human CD8^+^ T cells in fresh RPMI supplemented with 30 U/mL IL-2. Fresh media was added every 4 days.

### mtDNA analysis

Cells were harvested in 350 μL RLT lysis buffer and mixed in a volume of phenol/chloroform/isoamyl alcohol (25:4:1) (PCIAA). After mixing, the samples were centrifuged (16000g for 5 min), and 0.25–0.3 mL of the supernatant was mixed with 50 μg/μL glycoblue (AM9515, ThermoFisher), 0.3 M sodium acetate and 0.7x volume of isopropanol. After spinning samples at 16000g for 5 min at 4°C, DNA pellets were washed twice through addition of ice-cold 70% ethanol and centrifugation at 16000g for 10 min at 4°C. Air dried pellets (5-20 min) were then dissolved in 0.4 ml Tris–EDTA (TE) buffer. DNA was set to 2.5 ng/ml and analysed by qPCR using primers for nuclear 18S (FW 5’-TAGAGGGACAAGTGGCGTTC-3’, RV 5’-CGCTGAGCCAGTCAGTGT-3’) (Thyagarajan et al., 2013) and for mitochondrial DNA (FW 5’-GCCTTCCCCCGTAAATGATA-3’, RV 5’TTATGCGATTACCGGGCTCT-3’) (Venegas and Halberg, 2011).

### Single-Cell Calcium Imaging

Frozen splenocytes from donor TCR-transgenic OT-I mice were acclimatised for 3 h to either 21% or 1% O_2_ before activation with 1 μg/mL SIINFEKL peptide for 48h. 24 h post-activation, cells at hypoxia were moved to 21% and all cells were subsequently cultured at normoxia until d7. OT-I were kept in RPMI-1640 medium containing 2 mM glutamine, 10% FBS, 1% penicillin-streptomycin, 55 μM β-mercaptoethanol, supplemented with human IL-2 (50 U/mL 0-48h, 20 U/mL 48h+). At day 7, OT-I T-cells (∼400,000 in 1 mL growth media) were loaded with 1 μM Fura-2 AM (F1221, ThermoFisher) for 40 min RT, in dark, on a horizontal rocker. Cells were washed and resuspended in Ringer’s buffer (45 mM NaCl, 4 mM KCl, 10 mM Glucose, 10 mM HEPES, 2 mM MgCl2, 1 mM CaCl2, pH 7.4) plus 20 U IL-2, before being left to de-esterify Fura-2 and adhere to poly-ornithine coated coverslips for at least 20 min. Coverslips were washed once more and mounted with 300 μL Ringer’s solution + IL-2 on the microscopy set up (Zeiss Axiovert S100TV equipped with a pE-340fura (CoolLED) LED light source with LED 340 nm (excitation filter: 340/20) and 380 nm (excitation filter: 380/20) together with a T400 LP dichroic mirror and 515/80 emission filter, a sCMOS pco.edge camera and a Fluar ×20/0.75 objective). Ratiometric single-cell time-lapse imaging was performed at 5 s intervals with baseline fluorescence measured for 4 min, followed by another 6 min reading after addition of SIINFEKL (final concentration 4 μg/mL). VisiView 4.2.0.0 software (Visitron Systems) was used for data analysis and calculated 340 nm/380 nm fluorescence ratios (F_340nm_/F_380nm_) were taken as directly proportional to cytosolic [Ca^2+^].

### ELISA

To detect IFN-γ and IL-2 production, enzyme linked immunosorbent assays (ELISA) were performed on T cell conditioned media. For this purpose, Human IFN gamma Uncoated ELISA kit (88-7316, Invitrogen) and IL-2 Human Uncoated ELISA Kit (88-7025, Invitrogen) were used according to the manufacturer’s protocol and the signal was obtained in a microplate reader (Sunrise, Tecan Austria GmbH) at a wavelength of 450 nm.

### Statistics

Statistical analyses were performed with Prism 9 software (GraphPad). A P value of <0.05 was considered significant and the statistical tests and sample sizes (n) used are stated in figure legends. Normality tests and P values are provided in Source data files.

### Source data

Excel files are provided for access to all raw data and uncropped blot images. Data from Figures 1-5 was included in Source Data files 1-5, respectively.

## Acknowledgments

The authors gratefully acknowledge the flow cytometry facility from the Scholl of the Biological Sciences for their support and assistance in this work. Bioenergetic experiments were performed at the Medical Research Council Toxicology Unit, University of Cambridge, Cambridge, UK. The authors would like to thank Dr. Xiao-Ming Sun and Prof. Marion MacFarlane for providing access to the Seahorse XF96 Bioanalyser and helpful discussions. The work was funded by the Knut and Alice Wallenberg Scholar Award, the Swedish Medical Research Council (Vetenskapsrådet 2019-01485), the Swedish Cancer Fund (Cancerfonden, CAN2018/808), the Swedish Children’s Cancer Fund (Barncancerfonden PR2020-007), the Portuguese Foundation for Science and Technology scholarship to Pedro P. Cunha (SFRH/BD/115612/2016), a Canadian Institutes of Health Research Fellowship to Brennan J. Wadsworth and the Principal Research Fellowship (214283/Z/18/Z) to Randall S. Johnson from the Wellcome Trust.

## Author contributions

Conceptualization, P.P.C, E.M and P.V; Methodology, P.P.C, L.C.M.K and P.V; Investigation, P.P.C, E.M, L.C.M.K, R.M.H, D.B, B.J.W, L.B and C.B; Writing - Original draft, P.P.C; Writing - Review & Editing, P.P.C, E.M, L.C.M.K, R.M.H, B.J.W, I.P.F, I.B, P.V and R.S.J; Funding acquisition, R.S.J

## Supplementary figures

**Figure S1.**
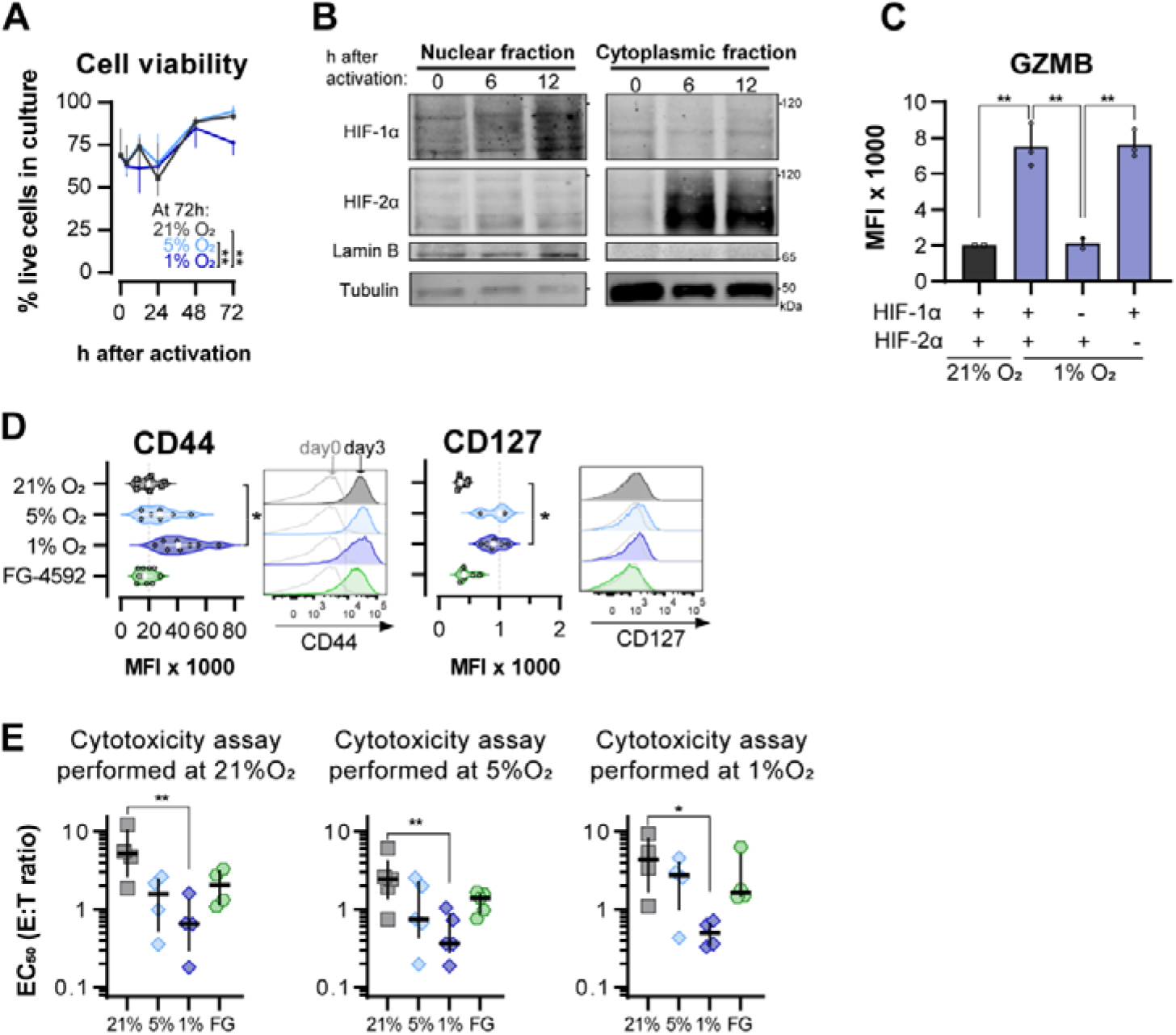
Hypoxia and inhibition of negative HIF signalling in CD8^+^ T cell effector differentiation (continued). **(A)** Proportion of viable of CD8^+^ T cells in cultures grown in 21%, 5% or 1% O_2_ determined with automated cell counter and trypan blue; n=4. **(B)** HIF-1α and HIF-2α protein expression in nuclear extracts from mouse CD8^+^ T cells activated in 21% O_2_. Lamin B and Tubulin were used as loading control for nuclear and cytoplasmic fractions, respectively. HIF-1α and Lamin B blotted with 800CW secondary antibodies and HIF-2α and Tubulin blotted with 680RD secondary antibodies. VHL knock-out T cells were used as positive control; n=1. **(C)** MFI of GZMB in WT (+) and *Hif1*^fl/fl^dLck^CRE^ *Hif2*^fl/fl^dLck^CRE^ cells (HIF-1α or HIF-2α KO, -) mouse CD8^+^ T cells cultures for 7 days in 21% (gray) or 1% O_2_ (blue); n=2-3. **(D)** OT-I CD8^+^ T cell differentiation was analysed by flow cytometry 3 days after activation. Histograms are representative flow cytometry plots for each parameter and are pre-gated on live, singlet, CD8^+^ events. The vertical dotted line defines in graphs the median of 21%-grown cells and in histograms the gate for each marker. Grey lines in the histograms represent the expression of these markers at day 0 prior to activation (naive). n=3-10. **(E)** *In vitro* cytotoxicity assay using the following conditions: Wild-type OT-I cells cultured in 21%, 5% or 1% O_2_; WT cells activated in 21% O_2_ and treated with 50 μM FG-4592. OT-I cells were co-cultured with B16-OVA cancer cells for at least 14 hours. EC_50_ values obtained from non-linear regression of dose-response curves obtained from co-cultures with different effector:target (E:T) ratios performed at 21% (left), 5% (middle) or 1% (right); Each data point represents and independent animal; n=4-5. All results shown as median ± IQR. * P<0.05, ** P<0.01, *** P<0.001; two-way ANOVA corrected with Tukey’s test (A) or Holm-Šídák’s multiple comparisons test (C-E). Full data and statistical analysis from Figure S1 can be found in Source data 1.

**Figure S2.**
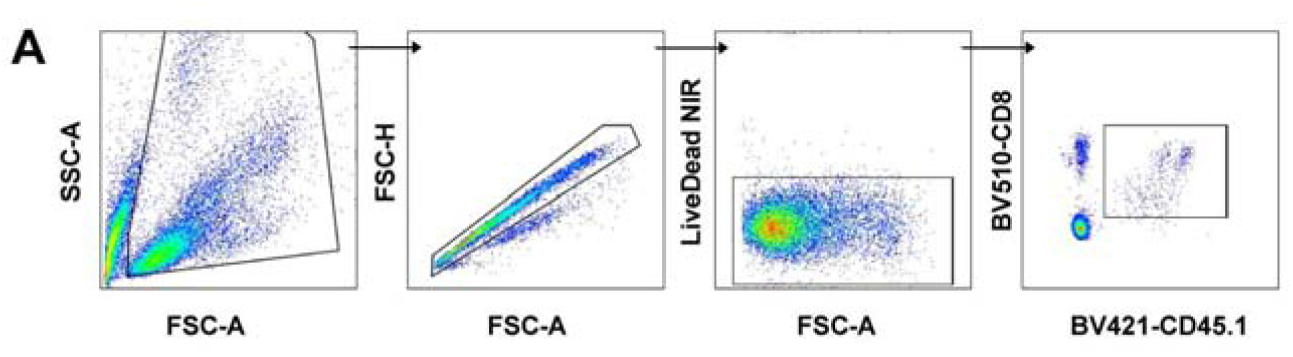
Effects of hypoxia and increased HIF signaling in antitumour function of CD8^+^ T cells (continued). **(A)** Gating strategy for analysis of peripheral blood.

**Figure S3.**
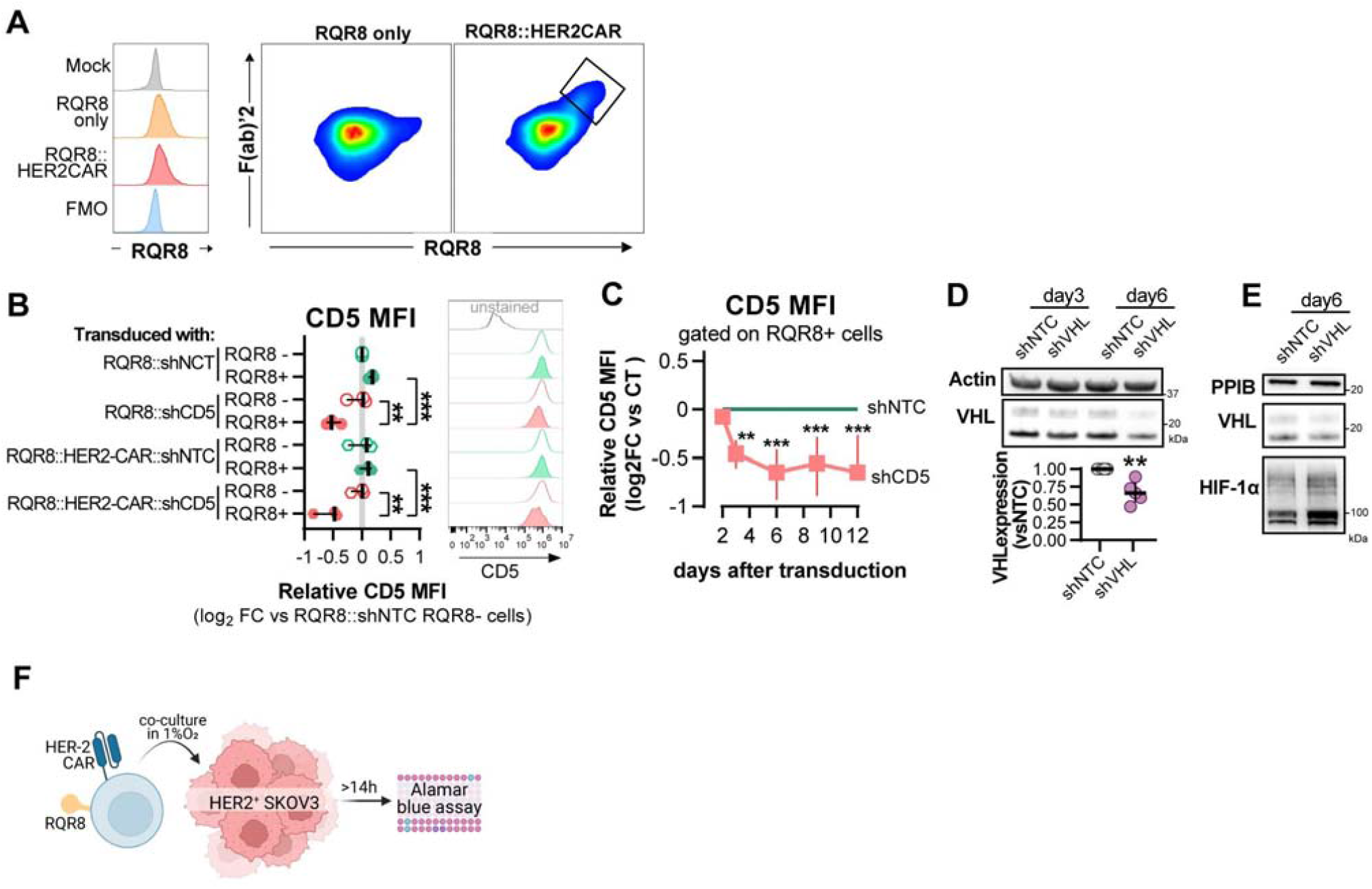
Validation of CAR and shRNA-CAR vectors. **(A)** Histograms of RQR8 staining with Qbend/10 CDE34 antibody (left) and correlation between RQR8 and surface CAR staining with F(ab)’2. **(B)** CD8^+^ T cells were transduced with a shRNA vector against non-target control (NTC) or CD5, alone or in combination with HER2CAR. On day 6, CD5 MFI was determined by flow cytometry in RQR8-(non transduced) and RQR8 (transduced) events. MFI values are presented as log 2 fold change (FC) relative to RQR8-cells from the population of cells transduced with RQR8-shRNA vectors. Histograms are representative flow cytometry plots for CD5 and are pre-gated on live, singlet, TCRαβ+ events. The dotted line in histograms was set according to the fluorescence peak of the control condition. Each data point represents an independent donor; n=3. **(C)** Activated human CD8^+^ T cells were transduced with a shRNA vector against non-target control (NTC) or CD5. MFI values of CD5 were analysed on days 2, 3, 6, 9 and 12 post-transduction and are presented as log2 FC relative to the NTC control in each day; n=3. **(D)** Western blot analysis of VHL protein in CAR-T cells 3 and 6 days following transduction with shNTC or shVHL. Actin was used as loading control. Top: representative immunoblot; bottom: quantification of VHL expression relative to shNTC on day 6 (left). **(E)** VHL and HIF-1a expression 6 days after transduction with shNTC or shVHL. **(F)** Cytotoxicity assay with HER2 CAR-T cells. CAR-T cell number (RQR8+) was determined by flow cytometry and cells were co-cultured with SKOV3 cell line in different effector:target rations for at least 14 hours in 1% O_2_. All results are shown as median ± IQR; **P<0.01, ***P<0.001; Two way ANOVA mixed effect paired analysis with Sidak multiple comparison correction (B) and Wilcoxon matched-pairs signed rank test relative to control (C-D). Full data and statistical analysis from Figure S3 can be found in Source data 3.

**Figure S4.**
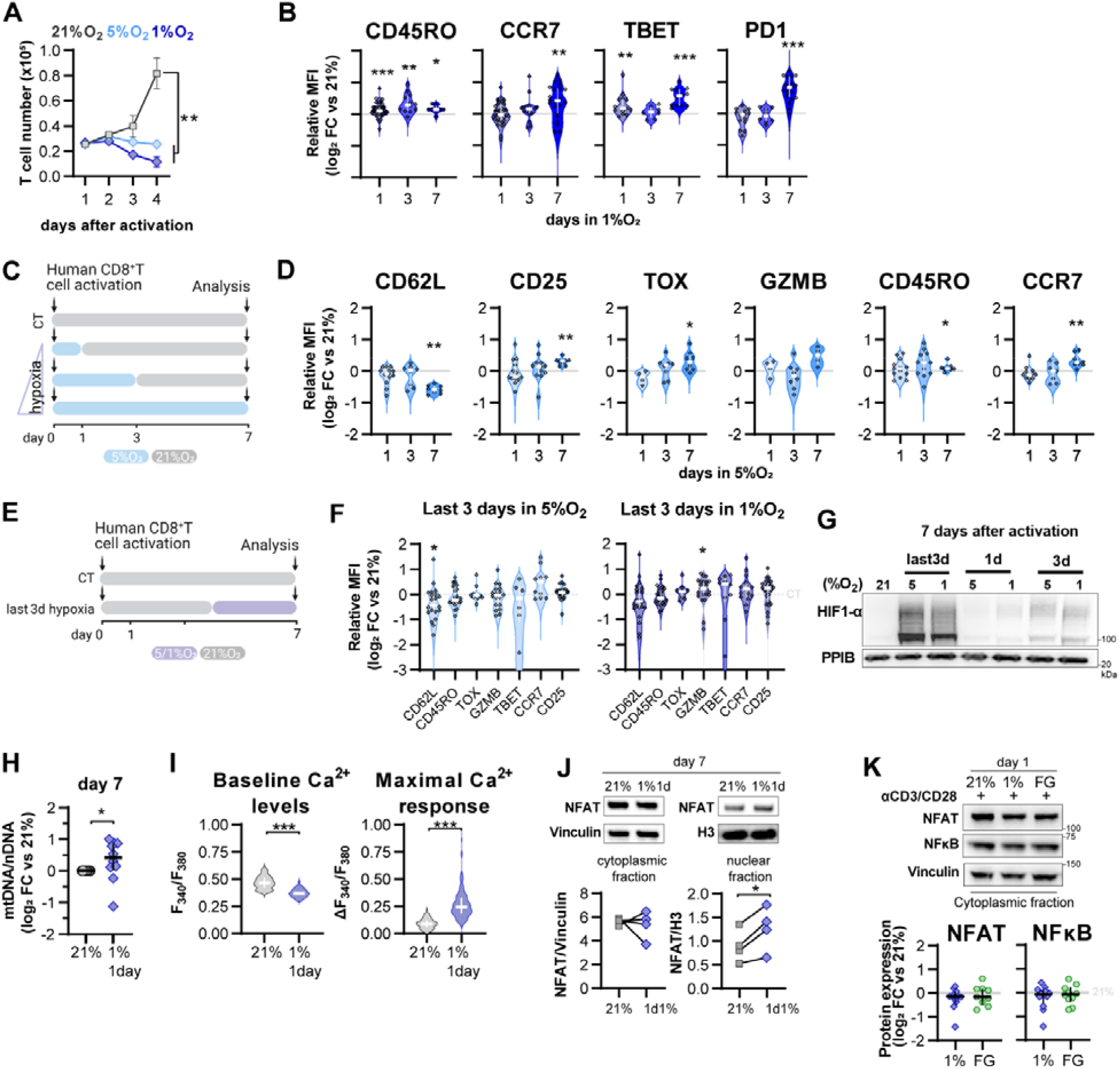
Effect of different *in* low oxygen conditioning protocols in human CD8^+^ T cell differentiation. **(A)** Expansion of human T cells in 21, 5 or 1%O2 as determined with automated cell counter; n=4. **(B)** Expression of differentiation markers determined by flow cytometry and shown as log2 fold change in MFI relative to 21% CT (horizontal grey line) following conditioning to 1% O_2_ as described in Figure 4A; n=8-38. **(C)** Human CD8^+^ T cells were activated in 5 %O_2_ for 1, 3 or 7 days. Cells continuously grown in 21% were used as control (CT). **(D)** Expression of differentiation markers determined by flow cytometry and shown as log2 fold change in MFI relative to 21% CT (horizontal grey line) following conditioning to 5% O_2_ as described in C; n=4-22. **(E)** Human CD8^+^ T cells were activated in 5 or 1% O_2_ for the last 3 days of a 7-day culture. Cells continuously grown in 21% were used as control (CT). **(F)** Expression of differentiation markers determined by flow cytometry and shown as log2 fold change in MFI relative to 21% CT (horizontal grey line) following conditioning to 5% (left) or 1% O_2_ (right) as described in E. **(G)** HIF-1α protein expression in nuclear extracts from human CD8^+^ T cells activated for 7 days. Conditions analysed: continuous culture in 21% O2 (CT); exposure for the last 3 days to 5 or 1% O2 (as shown in D); culture for 1 or 3 days in 5 or 1% as shown in B or Figure 4A, respectively. PPIB used as loading control. Representative immunoblot (n=3). **(H)** Mitochondrial DNA (mtDNA) content in CD8^+^ T cells activated in 1 %O_2_ for one day and changed to 21 %O_2_ for 6 days. Data shown in log2 fold change relative to cells continuously maintained in 21% for 7 days. Nuclear DNA (nDNA) was used to normalize mtDNA content; n=14.. **(I)** Baseline cytosolic Ca^2+^ level (left) and maximal Ca^2+^ response (SOCE) following addition of SINFEKL (right) in OT-I T-cells activated in 1% O_2_ for 1 day and expanded for 6 days in 21% O_2_. Data shown as violin plots with median ± IQR represented in white. Number of cells analysed (21% grey, n=249; 1%1day blue, n=128) are derived from multiple coverslips and 2 OT-I donor spleens. **(J)** NFAT protein expression in cytoplasmic and nuclear fraction of human CD8^+^ T cells activated for 24 h in 1% O_2_ and expanded for 6 days in 21% O_2_. Vinculin and histone 3 were used as loading controls for cytoplasmic and nuclear fractions, respectively. Representative immunoblots (top) and normalized protein expression (bottom); n=4. **(K)** Cytoplasmic protein analysis 24h post activation of human CD8^+^ T cells with anti-CD3/CD28 dynabeads in 21% O_2_, 1% O_2_ or 21% O_2_ + 50μM FG-4592 (FG). Vinculin was used as loading control. Representative immunoblots (left) and log2 FC of NFAT1 (middle) and NFkB (right) of cells activated in 1% or with FG relative to 21% controls; n=3-10. All results were pooled from at least two independent experiments and are shown as median ± IQR; * P<0.05, ** P<0.01, *** P<0.001; two-way ANOVA corrected with Tukey’s test (A) and Wilcoxon matched-pairs signed rank test relative to control (B-E). Full data and statistical analysis from Figure S4 can be found in Source data 4.

**Figure S5.**
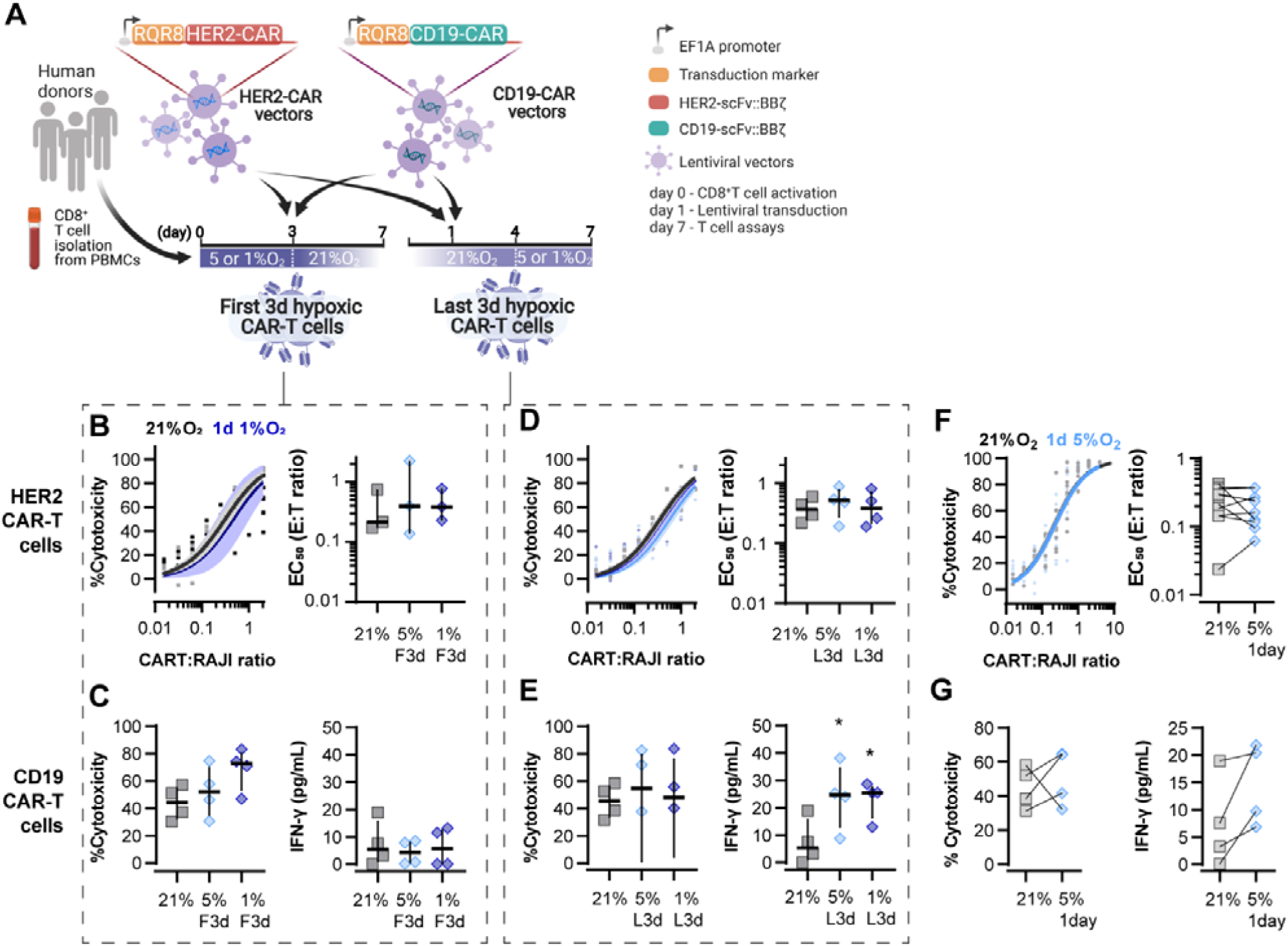
Evaluation of different protocols of low oxygen conditioning in CAR-T cell function. **A)** Human CD8^+^ T cells were activated, transduced with HER2- or CD19-CAR vectors the day after and assayed on day 7. Cells were activated in 5 or 1% O_2_ for 3 days amd swapped to 21% until day 7 or activated in 21% for 4 days and swapped to 5 or 1% O_2_ until day 7. CAR-T cells continuously maintained in 21% O_2_ were used as controls and RQR8 was used as transduction marker. **(B)** *In vitro* cytotoxicity assay with HER2 CAR-T cells conditioned to low oxygen for the first 3 days (F3d) of a 7 day culture as shown in A. HER-2 CAR-T cells were co-cultured with SKOV3 tumour cells at different effector:target (E:T) ratios and cytotoxicity was assessed after 14-18 hours of co-culture at 1% O_2_. Left: dose-response curves (plotted with 95% confidence intervals represented in shaded areas) determined with non-linear regression ([agonist] vs normalised response); right: EC_50_ values obtained from non-linear regression; n=4. **(C)** *In vitro* cytotoxicity assay with CD19 CAR-T cells conditioned to low oxygen for the first 3 days (F3d) of a 7 day culture as shown in A. CD19 CAR-T cells were co-cultured with RAJI tumour cells at a E:T ratio of 2:1 and cytotoxicity was assessed after 14-18 hours of co-culture at 5% O_2_. Left: cytotoxicity relative to cells transduced with vector control containing RQR8 alone; right: IFN-γ ELISA using supernatants of the co-culture between CAR-T cells and RAJI; n=3-4. **(D)** *In vitro* cytotoxicity assay with HER2 CAR-T cells conditioned to low oxygen for the last 3 days (L3d) of a 7 day culture as shown in A. HER-2 CAR-T cells used as described in B; n=4. **(E)** *In vitro* cytotoxicity assay with CD19-CAR T cells conditioned to low oxygen for the last 3 days (L3d) of a 7 day culture as shown in D. CD19 -2 CAR-T cells used as described in C; n=3-4. **(F)** Same experiment described in Figure 5B but with conditioning to 5% O_2_ instead of 1%O_2_. **(G)** Same experiment described in Figure 5C but with conditioning to 5% O_2_ instead of 1%O_2_. All results shown as median ± IQR; * P<0.05, Wilcoxon matched-pairs signed rank test relative to control. Full data and statistical analysis from Figure S1 can be found in Source data 5.

## Supplementary tables

**Table S1.**
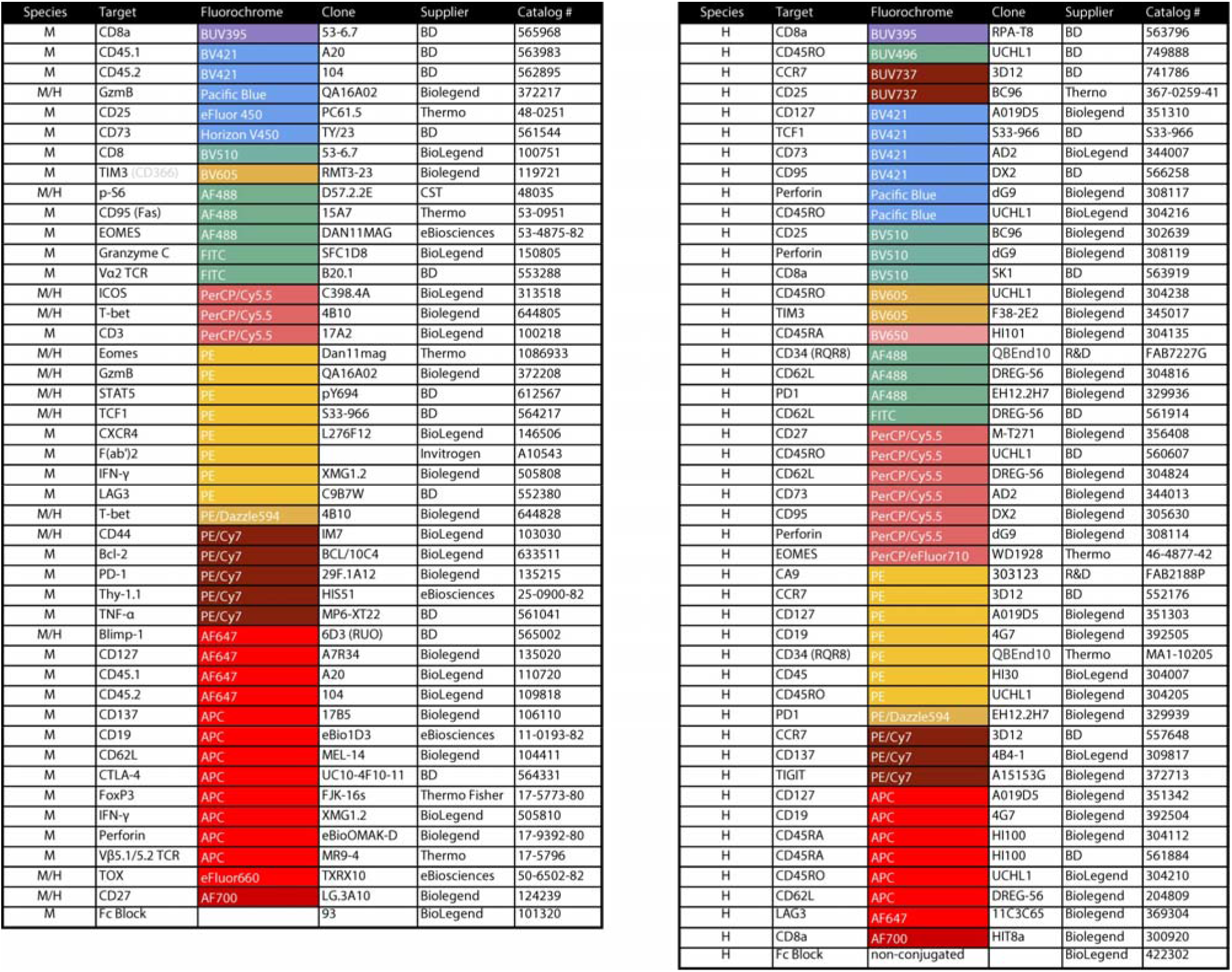
List of antibodies used for flow cytometry. Species reactivity: mouse (M), human (H) or both (M/H).

**Table S2.** List of antibodies used for western blot analysis.

| Target | Catalog Number | Supplier |
| --- | --- | --- |
| GLUT1 | ab115730 | abcam |
| Vinculin | ab219649 | abcam |
| HDAC | ab156064 | abcam |
| HIF-1α | 610958 | BD Biosciences |
| PPIB | 43603 | Cell Signaling Techonology |
| PPIB | 43603 | Cell Signaling Techonology |
| PGC1α | 4259 | Cell Signaling Techonology |
| NFIL3 | 14312 | Cell Signaling Techonology |
| NFAT1 | 4389 | Cell Signaling Techonology |
| NF-κB | 8242 | Cell Signaling Techonology |
| cMYC | 5605 | Cell Signaling Techonology |
| LDHA | 3582 | Cell Signaling Techonology |
| Lamin b1 | 13435 | Cell Signaling Techonology |
| HIF-1α | NB-100-449 | Novus |
| HIF-2α | AF2997 | R&D Systems |

**Table S3.**
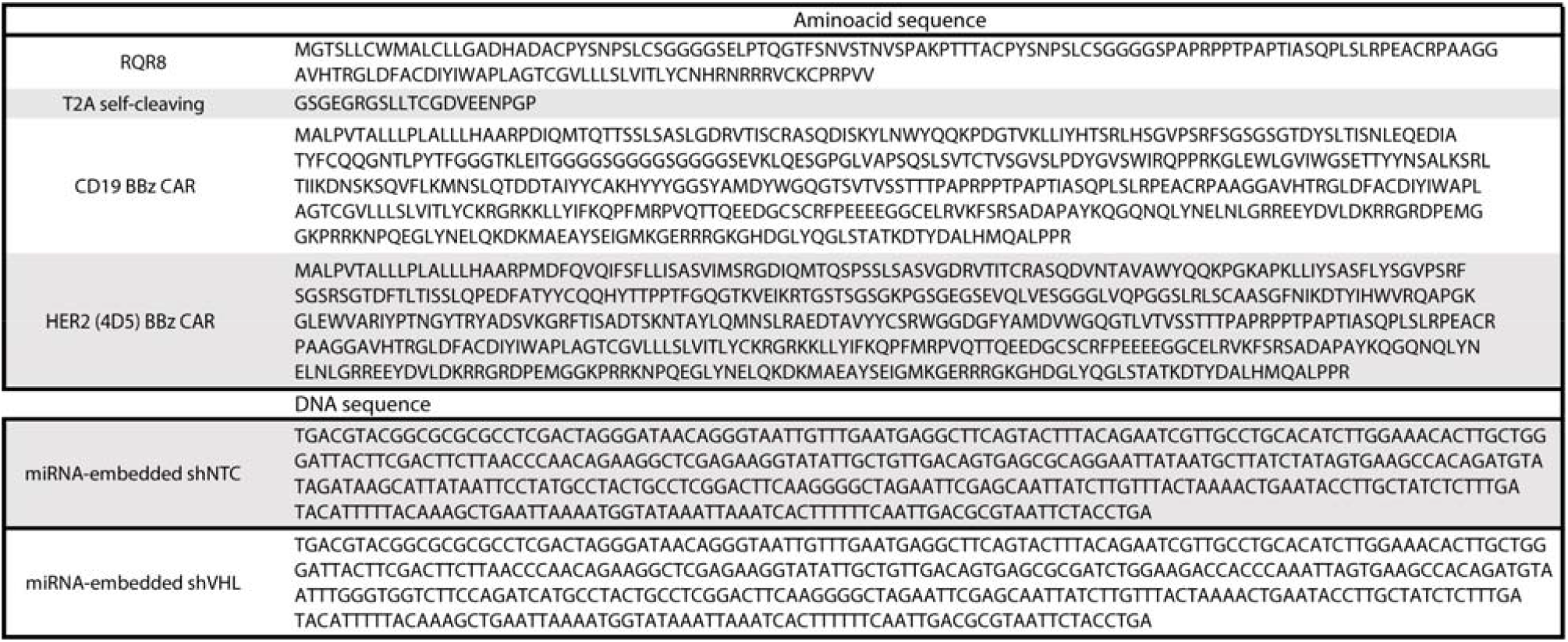
List of protein sequences and DNA sequences corresponding to miRNA-embedded shRNAs used in viral vectors.

